# Temporal phosphoproteomic programmes set a redox-gated rheostat on neutrophil phagocytosis

**DOI:** 10.64898/2026.10.26.761950

**Authors:** Maire Röder, Muhammad Iqbal, Anthony YKC Ng, Helena Brown, Iat Tai Wong, Samuel Lewirt, Motaharehsadat Heydarian, Ashley D Smith, Isabella M Monsanto, Mengye Li, Alexander JT Wood, Matthew Routledge, Kamal Kishore, Clive D’Santos, Eva Papachristou, David C Thomas, Klaus Okkenhaug, Andrew Conway Morris

## Abstract

Neutrophils are essential frontline immune cells in the defence against bacterial pathogens such as *Staphylococcus aureus*, but their killing machinery also causes host tissue damage. How neutrophils regulate these processes remains incompletely understood. Here, we profiled the proteome and phosphoproteome of human neutrophils across a two-hour exposure to *S. aureus*, alongside five simultaneous functional readouts, before investigating how an initial encounter modulates the response to a second. The phosphoproteomic response resolved into qualitatively distinct early and late windows: the early landscape drives particle uptake, while by the late window this machinery is deactivated, with a broadly supressed kinome, and cells geared towards inflammatory output. Correlating protein phosphorylation and functional kinetics — an approach we term KINETIC — implicates putative mediators of key neutrophil antimicrobial functions. A genome-wide CRISPR knockout screen in neutrophil-like PLB-985 cells, sorted on phagocytic load, recovered canonical engulfment machinery among the strongest facilitators of phagocytosis, additionally identifying the lysosomal Ragulator-mTORC1 nutrient sensing pathway and HOPS complex corresponding to the KINETIC-identified pathways. Conversely, the NRF2 repressor KEAP1 was the strongest phagocytic brake, positioning oxidative burden as a regulator of phagocytic appetite. Neutrophils in the late window exhibited impaired phagocytosis and phagosomal acidification on *S.aureus* re-challenge. Underlying these functional deficits, cells in the late window were refractory across multiomic layers: a second *S. aureus* meal altered no phosphoprotein, protein, or transcript, whereas the chemoattractants PAF and fMLP evoked a largely consistent phosphoproteomic programme, delineating the phenotype as receptor-specific rather than a global loss of signalling capacity. Consistent with de-repressing the antioxidant response via genetic loss of KEAP1 increasing phagocytosis, reducing ROS production via inhibition of the NADPH oxidase resulted in a partial rescue of phagocytosis of the re-exposed *S.aureus*. Together, these findings describe a ROS-gated phagocytic rheostat, enabling neutrophils to meter their phagocytic appetite according to their exposure history, likely limiting uptake of pathogens they lack the capacity to effectively digest. Inappropriate activation of this rheostat under conditions of physiological stress or severe illness may underpin the defects in phagocytosis identified in such patients.

## 1 Introduction

Neutrophils are the most abundant leukocyte in the human circulation and are rapidly recruited to sites of infection and inflammation. High-throughput single-cell technologies have redefined neutrophils as a dynamic and heterogeneous population, where functional plasticity, rather than the existence of stable subpopulations alone, underpins their adaptability to signals such as cytokines, pathogens, and metabolites [1, 2]. Absence or dysfunction of neutrophils markedly increases susceptibility to infection [3, 4] but neutrophils are also key mediators of tissue damage in sepsis [5, 6] and drive organ failure and increased mortality. Impaired neutrophil antimicrobial functions and neutrophil-mediated tissue damage occur simultaneously in sepsis [3, 7, 8], and the mechanisms that underpin these divergent maladaptive states remain incompletely understood.

Phagocytosis and phagosomal maturation are central to bacterial killing [9], and involve acidification by proton pumps, fusion of granules containing proteolytic enzymes, and the production of reactive oxygen species (ROS). Failure of this maturation process can lead to intracellular persistence of the engulfed pathogen, allowing the neutrophil to act as a ‘Trojan horse’ and seed infection elsewhere [10–12]. Although key signalling pathways have been identified in these processes [9], the networks that orchestrate the precise sequence of events and any safeguards that mitigate against intracellular pathogen persistence are poorly characterised, representing a question of central biomedical importance in our understanding of infection.

Neutrophils are notable for their exuberant production of ROS, primarily via the abundant NADPH oxidase NOX2, a multisubunit complex comprising two transmembrane (gp91*^phox^* /NOX2 and p22*^phox^*) and three cytosolic (p47*^phox^*, p67*^phox^*, and p40*^phox^*) subunits that interact with the small G-proteins RAC1 or RAC2 [13]. NOX2 is located on the plasma and phagosomal membranes, facilitating extracellular ROS release and intra-phagosomal oxidative killing, respectively. It is increasingly clear that ROS in neutrophils not only have important bactericidal effects, but also subsume signalling roles. ROS play a central role in the generation of neutrophil extracellular traps (NETs), where activation of the RAF-MAPK-ERK [14] and direct oxidation of DNA [15] appear to be central. However, ROS-mediated signalling has also been implicated in limiting neutrophil tissue infiltration [16] and pathogen ingestion [17]. The cellular response to this oxidative burden is itself regulated, principally via the NRF2 axis. Furthermore, ROS are capable of modulating protein function post-translationally via reversible oxidation of cysteine residues [18], a process with significant immunomodulatory precedent: oxidation of interferon regulatory factor 1 (IRF1) Cys83 restrains type 1 interferon signalling by reducing nuclear translocation and transcriptional activity [19].

Previous work in macrophages and monocytes revealed that the response of these phagocytes to stimuli is shaped by their encounter history: trained immunity heightens subsequent responses whereas endotoxin tolerance selectively suppresses them. Both phenomena are mediated cell-intrinsically by epigenetic and metabolic reprogramming. In neutrophils, which are shorter-lived, modulation of innate responses has thus far been characterised at the level of granulopoeisis, where reprogrammed progenitors alter the character of newly generated neutrophils over a longer timeframe. Whether mature neutrophils already in circulation or tissue spaces adjust their own behaviour according to what has already been ingested, is not known [20–24].

Previously, we developed a neutrophil phosphoproteomic profiling system that enabled the discovery of novel features of C5a-mediated phagocytic dysfunction [25]. There, we revealed striking changes in the neutrophil phosphoproteome in response to *Staphylococcus aureus*, but this was a single timepoint study. Here, we expand on these findings, leveraging longitudinal phosphoproteomics to deconvolute discrete neutrophil effector functions during *S. aureus* challenge. Following the identification of distinct early and late profiles, we investigated the matching of functional and phosphoproteomic kinetics using a technique we term ‘Kinase Network Inference through Trajectory Integration and Correlation (KINETIC)’ to identify novel interactors across key bactericidal processes, and observe markedly reduced activity across a broad range of kinases following prolonged pathogen exposure. We then examined the effects of prior Staphylococcal exposure on subsequent responses to re-challenge in these windows, and how these differed across the transcriptional, protein, and post-translational modification landscapes. Our investigations reveal a phenotypic shift in neutrophils following prolonged pathogen exposure, exhibiting reduced phagocytosis and phagosomal acidification but transcriptional and translational evidence of pro-inflammatory signalling and chemokine production. We identify a putative ROS-sensitive phagocytic rheostat that limits phagocytosis during serial pathogen exposure. These findings provide novel insights into the regulatory networks that control neutrophil responses to pathogens, and identify a ROS-dependent phenotype induced by prolonged pathogen exposure that has implications for our understanding of pathogen control and immunopathology. Importantly, a mechanism equipping mature neutrophils with the ability to rapidly and durably modulate downstream functionality in response to phagocytic history represents a novel paradigm. Not only would this constitute a safeguard mitigating uptake of pathogens that cannot be effectively neutralised, but it may become maladaptive in severe infection, constituting a mechanistic explanation for the paradox of sepsis, and opening therapeutic avenues to manipulating this.

## 2 Results

### 2.1 Exposure to *S. aureus* induces a dynamic, biphasic phosphoproteomic response

To characterise the neutrophil response to *S. aureus*, we performed quantitative phosphoproteomics and proteomics in neutrophils from healthy human donors. Cells were sampled immediately following purification, prior to stimulation (baseline), and over a seven-point time course (2 to 120 minutes) after exposure to heat-killed *S. aureus* bioparticles with time-matched controls at 30, 60 and 120 minutes (Figure 1A).

**Fig. 1.**
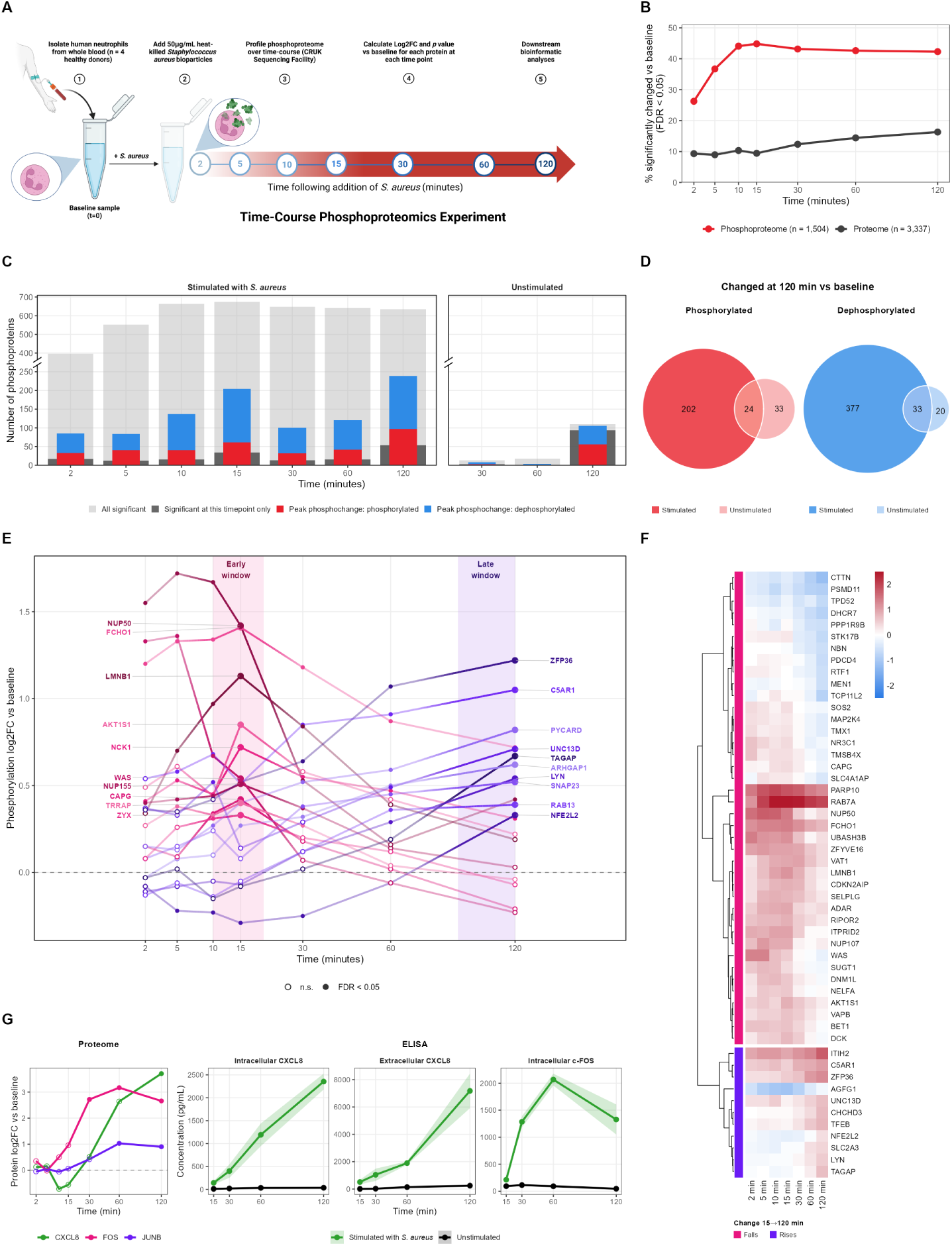
Exposure to *S. aureus* induced a dynamic, biphasic phosphoproteomic response. (A) Experimental design schematic: Neutrophils were isolated from whole blood healthy donors (=4), stimulated with 50 *µ*g/mL heat-killed, serum-opsonised *S. aureus* bioparticles, and sampled at 2, 5, 10, 15, 30, 60 and 120 minutes, with a pre-stimulation baseline and time-matched unstimulated controls at 30, 60 and 120 minutes. (B) Percentage of quantified features significantly changed from baseline (FDR *<* 0.05) at each timepoint, for the phosphoproteome (n = 1,504) and the neutrophil-intrinsic proteome (n = 3,337). (C) Number of phosphoproteins changed at each timepoint in stimulated (left) and unstimulated (right) cells. Grey, all significant; dark grey, significant at that timepoint only; red and blue, proteins reaching peak phosphorylation or dephosphorylation. (D) Overlap between phosphorylated and dephosphorylated phosphoproteins at 120 minutes in stimulated and unstimulated cells. (E) Phosphorylation trajectories (log_2_FC vs baseline) of representative early and late peaking phosphoproteins. (F) Phosphoproteins significant at 15 or 120 minutes (FDR *<* 0.05 at either) whose phosphorylation changed by more than 0.5 log_2_ units between the two windows, with falling and rising groups clustered separately. (G) Left, proteome trajectories of CXCL8, FOS and JUNB (log_2_ fold change versus baseline; filled points, FDR *<* 0.05). Right, ELISA validation in stimulated and unstimulated cells: intracellular CXCL8, extracellular CXCL8 and intracellular c-FOS.

Of 1,504 phosphoproteins quantified, 978 exhibited significantly different phosphorylation relative to baseline at one or more time-point (False discovery rate;FDR*<* 0.05). The response was overwhelmingly driven by stimulation rather than time *ex vivo*: unstimulated neutrophils showed minimal phosphoproteomic change over the first hour following purification (Figure 1C), whilst phosphoproteins changing after 120 minutes unstimulated ex-vivo showed limited overlap with those arising from pathogen stimulation (Figure 1D).

The proteome revealed a rapid rise in 123 plasma proteins at two minutes (Figure 1B), consistent with ingestion alongside opsonised bioparticles. To analyse the ’neutrophil intrinsic’ proteome and its change over time we excluded these plasma proteins, and used the remaining 3,337 proteins detected at the two-minute timepoint as the baseline dataset.

The phosphoproteome and proteome responded on timescales that exhibited little overlap. The proportion of the phosphoproteome carrying a significant change rose from 26.3% at 2 minutes to 44.8% by 15 minutes (Figure 1B) and then plateaued, whereas the proteome climbed gradually and had not plateaued at 120 minutes. Furthermore, the proteome exhibited a less substantial and dynamic response to *S. aureus* with fewer significant changes at each timepoint. Changes in phosphoprotein abundance did not relate to total protein abundance; of the 945 proteins quantified in both datasets, phosphorylation and abundance fold changes were not correlated at any timepoint.

### 2.2 Phosphorylation kinetics are bimodal, and peaks of phosphochange correspond to distinct early and late windows

Although the number of cumulative phosphorylation and dephosphorylation rose to a plateau relatively early in the timecourse, the timing of these changes was bimodal (Figure 1B) with peak phosphochange achieved at 15 minutes (204 proteins) and 120 minutes (239 proteins). Although dephosphorylation was more prevalent than phosphorylation overall, among strongly modulated proteins (log_2_ fold change; log_2_FC > 0.5) the balance inverted and then reversed with time: the phosphorylation:dephosphorylation ratio fell from 2.11 at 2 minutes to 0.66 at 120 minutes. These peaks of phosphochange denote qualitatively distinct early and late windows, suggesting that neutrophils shift between different functional states.

The early window is configured for ingestion. At 15 minutes, the phosphoproteome was dominated by the machinery of particle uptake [26]. The ITAM adaptor FCER1G and the PI3K regulatory subunit PIK3R2 peak in this window; PIK3R5 rose and plateaued, PIK3R1 is dephosphorylated and the RAC and CDC42 exchange factors and adaptors ARHGEF7, NCK1, DOCK2, DOCK5 and VAV3, all approached peak phosphorylation. Proteins with key roles in actin dynamics, including WAS, PAK2, WASF2, RCSD1, and CAPG, were significantly phosphorylated, as were the membrane-cytoskeleton adaptors DBNL and ZYX. Concurrently, there was a strong nuclear envelope modification signal: the nuclear pore complex components NUP50, NUP107, and NUP155, the pore-anchoring protein RANBP2, the nuclear basket protein TPR, and nuclear envelope component Lamin B1 were amongst the most strongly phosphorylated proteins. Lamin phosphorylation reduces nuclear rigidity [27]; the prevalence of nuclear envelope proteins alongside canonical mediators of pathogen engulfment may reflect the rapid nuclear deformation required during phagocytosis, alongside transcription factor shuttling. ERK/MAPK signalling was engaged within 2 minutes, with MAPK1 and MAPK3 phosphorylation peaked early and then returned to baseline over the timecourse. This was echoed in the gradual rise of RPS6KA2 phosphorylation to its peak at 30 minutes. The proteomic response was negligible at this point: only two proteins exhibited altered abundance at 10 minutes, rising to 43 by 15 minutes. The early window is thus reliant on post-translational reconfiguration of the proteins already present.

The late window consisted of a shift towards inflammation, degradation, secretion, and biosynthetic pathways. By 120 minutes, the pro-engulfment landscape seen at 15 minutes had largely disappeared and the landscape was shaped by signals that constrain further activation. Phosphorylation of NUP107, NUP153, NUP155, Lamin B1, WAS, and MAPK1/3 returned to baseline, whilst inhibitors of particle uptake accumulated phosphorylation: ARHGAP25, the Rac GTPase-activating protein, plateaued by 30 minutes. Similarly SHIP1 was stably phosphorylated throughout whilst phosphorylation of VSIR and SIRP*α* rose along with ARHGAP1 and ARHGAP45. LYN, an Src kinase with known pleiotropic roles in the coordination of phagocytosis and phagosomal acidification [28, 29], was also progressively phosphorylated.

Whilst particle uptake signals dropped out of the phosphoproteome over time, proteins involved in antimicrobial processing remained prevalent. Proteins involved in phagosomal maturation and acidification were sustained or continued to rise: RAB7A declined from its 15-minute peak but remained the most strongly phosphorylated protein in the dataset at 120 minutes (Figure 1E), alongside the v-ATPase subunit ATP6V1G1 and mRAVE complex member DMXL2, which assembles the v-ATPase proton pump [30]. The NADPH oxidase component NCF2 (p67*^phox^*) was increasingly phosphorylated over time, as were the exocytic machinery components VAMP8 and SNAP23, alongside the chemoattractant receptors C5AR1 and CXCR1. Meanwhile, the machinery required for protein synthesis gained phosphorylation progressively: RPS6KA2 and RPS6 phosphorylation peaked at 30 minutes whilst NCBP1 was significantly phosphorylated at 2 minutes and remained so thereafter. ZFP36, whose phosphorylation stabilises CXCL8 and TNF transcripts [31], became significant at 15 minutes and rose steeply to its maxima at 120 minutes. These findings tracked the proteome: FOS became significant by 30 minutes, JUNB by 60, and CXCL8 by 120 minutes. FOS and JUNB are AP-1 transcription factor subunits, and drive transcription of genes encoding inflammatory cytokines, chemokines, and growth factors [32]. Their abundance peaked at 60 minutes and declined in the late window. CXCL8 and FOS expression were also quantified by ELISA to validate (Figure 1G); both profiles being consistent with the proteomic data. The rise in intracellular CXCL8 preceded that of extracellular CXCL8, indicating denovo synthesis followed by secretion. Of the 125 proteins that changed abundance by 120 minutes, most (84) decreased. L-selectin abundance fell from 15 minutes, consistent with the shedding known to accompany neutrophil priming and activation [33]. The loss of FCGR3B (CD16) at 60 minutes is another characteristic of activation, whilst the loss of granule protein MMP9 indicates degranulation. Total integrin-alpha-M (CD11b) protein did not rise across the timecourse, nor did its heterodimer partner integrin beta-2(CD18), however surface CD11b rose (Supplementary Figure 1E). CD11b was also retained in its activated conformation (Supplementary Figure 1F), reflecting secretory granule mobilisation and activation . CXCR4, the canonical marker of neutrophil ageing, did not rise in stimulated cells even after 24 hours (Supplementary Figure 1C). We therefore infer that stimulated neutrophils in the late window not overtly de-primed nor aged, but have modulated their surface receptor repertoire.

The most variable features indicate a coordinated state transition. Of the proteins significant in both the proteome and phosphoproteome at 15 or 120 minutes, 51 had a difference in log_2_FC of at least 0.5 between the two time points (Figure 1F). Both these were asymmetric, with losses outnumbering gains (40 versus 11 phosphoproteins; 14 versus 4 proteins) and corroborate the shift in character of the phosphoproteome and proteome over time. Nuclear factor erythroid 2-related factor 2, a transcription factor that drives the production of anti-oxidant defences increased phosphorylation with time, placing the oxidative stress response among the small number of programmes that are switched on, rather than off, as the cell enters the late window. We hereafter refer to this transition as the phenotypic shift, and in subsequent experiments used the early and late windows to further interrogate it.

### 2.3 Successive waves of phosphorylation take the neutrophil from early to late phenotype

Although the biphasic pattern noted was the most prominent, interrogation provided little insight into how the proteins interact or subsume functional roles beyond those already well annotated in the literature. To interrogate the temporal architecture of phosphochange in more detail we used Dynamic Time Warping (DTW-SBD) to group proteins by the shape of their phosphorylation trajectories [34]. This resolved an ordered shift in the character of the response (Figure 2A). Each cluster contained an assembly of proteins with coherent functions (Figure 2B). We then overlaid STRING-DB protein-protein interactions within each cluster and ranked members by a composite score of intra-cluster connectivity, magnitude of phosphorylation change, and contribution to the cluster’s Gene Ontology enrichment (Table 1) to understand the interactions and functions subsumed by each cluster.

**Fig. 2.**
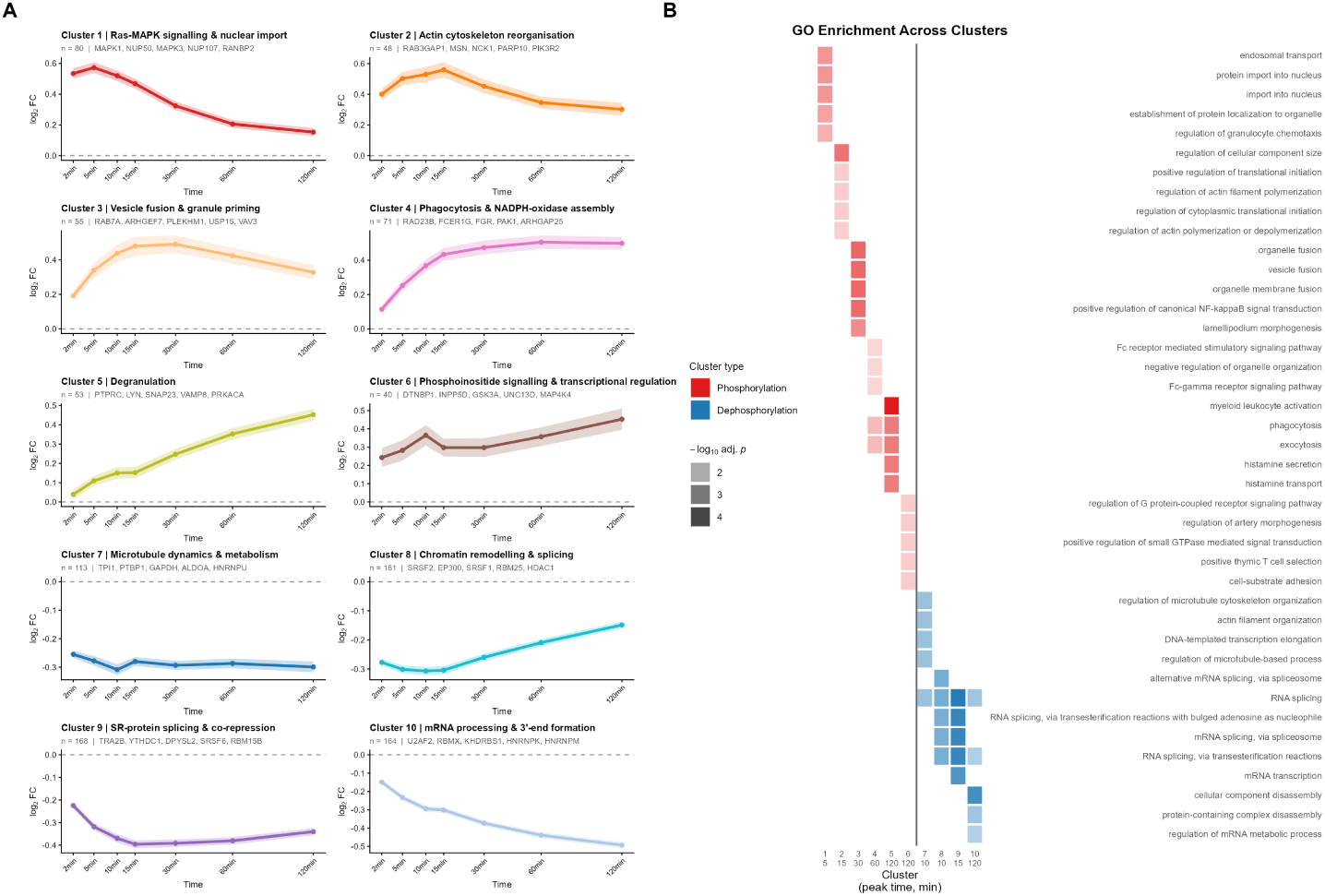
Successive waves of phosphorylation carry the neutrophil from the early to the late phenotype. (A) The ten shape-based clusters of phosphorylation trajectories derived by dynamic time warping, agnostic to function or absolute effect size. Each panel shows the cluster centroid and interquartile range, number of phosphoproteins belonging to each cluster, top hub proteins, and peak time. (B) Gene Ontology biological process over-representation analysis across clusters, computed against a whole-genome background with Benjamini-Hochberg correction within clusters.

**Table 1.**
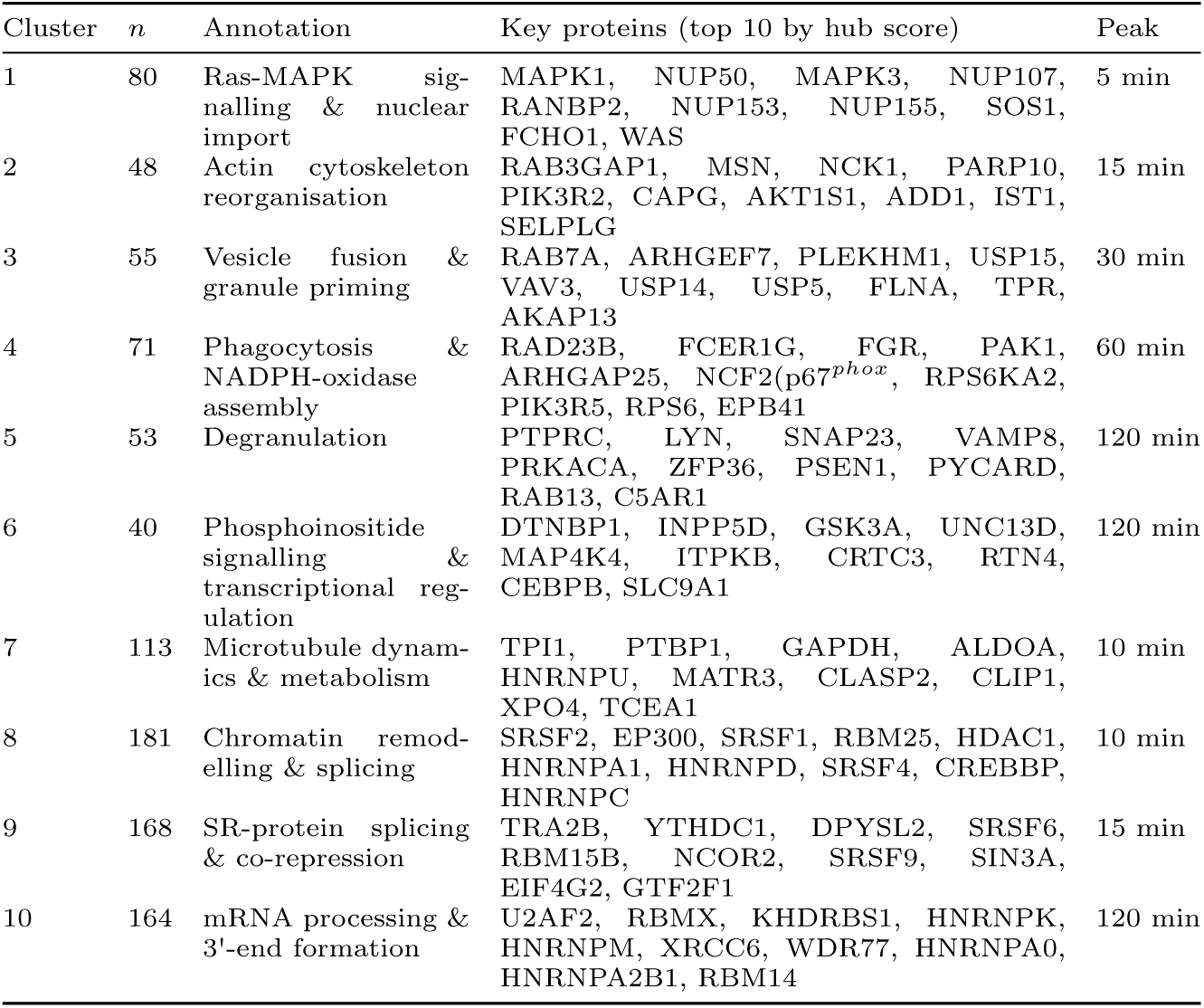
Shape-based clusters: functional annotations and top ten hub proteins. *n* is the number of phosphoproteins assigned to each cluster. Hub proteins are ranked within each cluster by a composite score combining intra-cluster STRING connectivity, magnitude of phosphorylation change and contribution to the cluster’s Gene Ontology enrichment (Methods). STRING interactions are taken at high confidence (combined score *≥* 700).

**Table 2.**
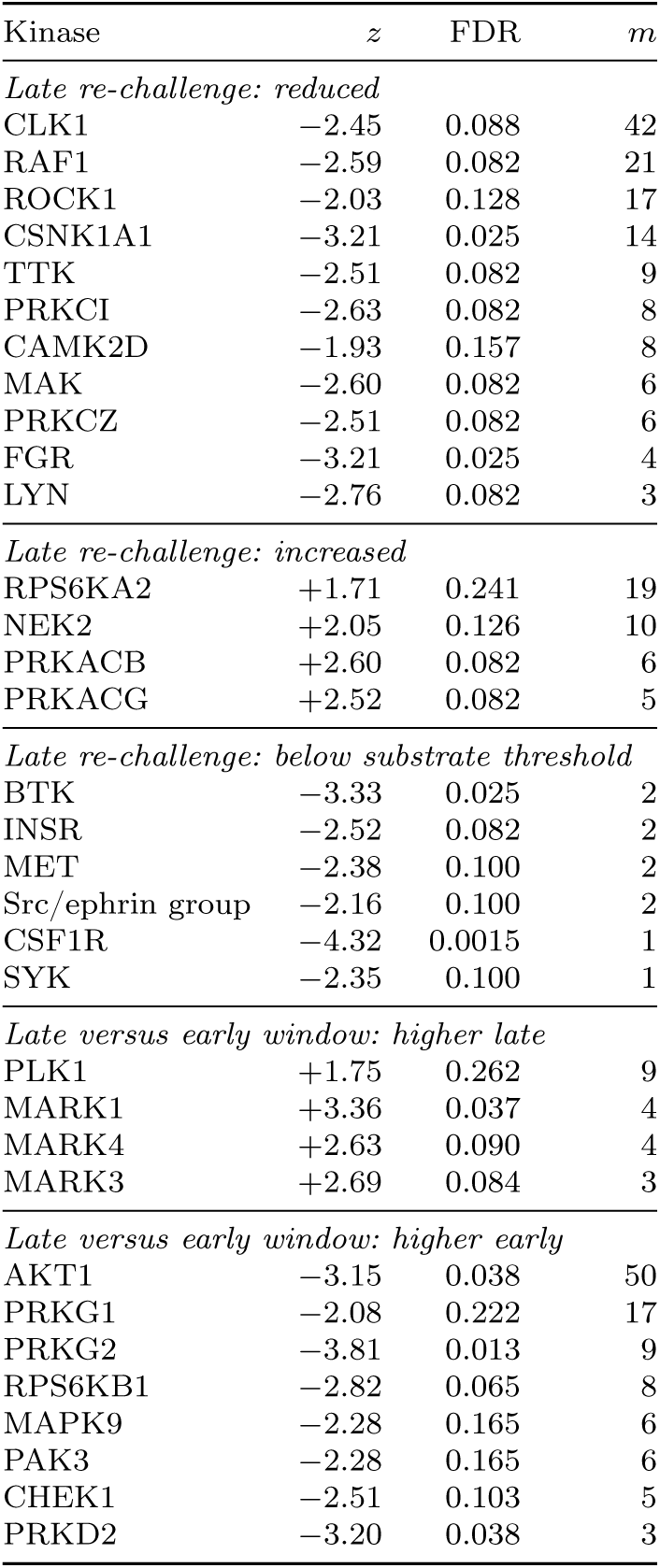
Kinases with altered inferred activity in the double-stimulation contrasts. Kinase–substrate enrichment analysis across 188 scored kinases; *z* is the enrichment score and *m* the number of annotated substrates supporting the inference. Within each block kinases are ordered by *m*.

Critically, this partitioning methodology is agnostic to protein function and absolute effect size [35] and recovered the early to late shift in character identified above. Among the six phosphorylated clusters, peak times moved through a coherent functional sequence: Ras-MAPK signalling and nuclear transport peaked at 5 minutes (MAPK1, MAPK3, NUP50, NUP107, RANBP2), actin remodelling at 15 minutes (MSN, NCK1, PIK3R2, CAPG), and vesicle fusion and granule priming at 30 minutes (RAB7A, ARHGEF7, PLEKHM1, VAV3). Phagocytosis and NADPH oxidase machinery (NCF2 (p67*^phox^*), FCER1G, FGR, PAK1, ARHGAP25) rose to a peak at 60 minutes and plateaued, whilst degranulation (PTPRC, LYN, SNAP23, VAMP8) and transcriptional regulators (CEBPB, CRTC3, GSK3A, INPP5D, ITPKB) continued to rise across the timecourse, peaking at 120 minutes. The early and late windows identified above therefore represent opposite ends of a continuous, ordered progression rather than discrete states.

#### Dephosphorylated clusters show different kinetics

Examination of the dephosphorylated clusters indicated that the late window did not simply reflect a decay of the early window. Microtubule and metabolic machinery (TPI1, GAPDH, ALDOA, CLASP2) and chromatin remodelling and splicing machinery (SRSF2, EP300. HDAC1, CREBBP) were dephosphorylated within the first 15 minutes and remained so, while mRNA processing and 3’-end formation machinery (U2AF2, HNRNPK, KHDRBS1, HNRNPM) was progressively dephosphorylated through to 120 minutes. This points to early and sustained dephosphorylation of metabolic and chromatin machinery accompanied by the activating cascade, whilst a second phase of dephosphorylation of mRNA-processing apparatus deepened into the late window.

This approach therefore refined the identity of early and late windows, and indicated how the progression emerges as the landscape moves from pro-engulfment to inflammatory and secretory.

### 2.4 Kinome Profiling Reveals a Broad Suppression of Kinase activity in the late Window

To identify the kinases underpinning the phenotypic shift, we used kinase-substrate enrichment analysis (KSEA) of the phosphoproteomic timecourse data, annotated by experimental (PhosphoSitePlus) and computational (NetworKIN) kinase asignments. We then sought to validate the dominant serine kinase findings using direct kinome profiling on a serine/threonine peptide array (PamChip STK).

Both approaches identified a striking reversal in kinase activity between the early and late windows. The number of KSEA-inferred active kinases fell from 18 at 15 minutes to 10 at 120 minutes, while those inferred as inhibited rose from 3 to 6 (Figure 3B). The contrast was starker in the kinome profiling array: of 90 kinases scored, 24 were significantly activated and none inhibited at 15 minutes, inverting to one activated and 27 inhibited at 120 minutes (Figure 3A). Across the kinases scored by both methods, 22/40 agreed in direction, and 7 reached significance in both. The activity of the RSK family rose in both methods in the early window and that of cGMP-dependent kinases fell in both methods in the late window (Figure 3C).

**Fig. 3.**
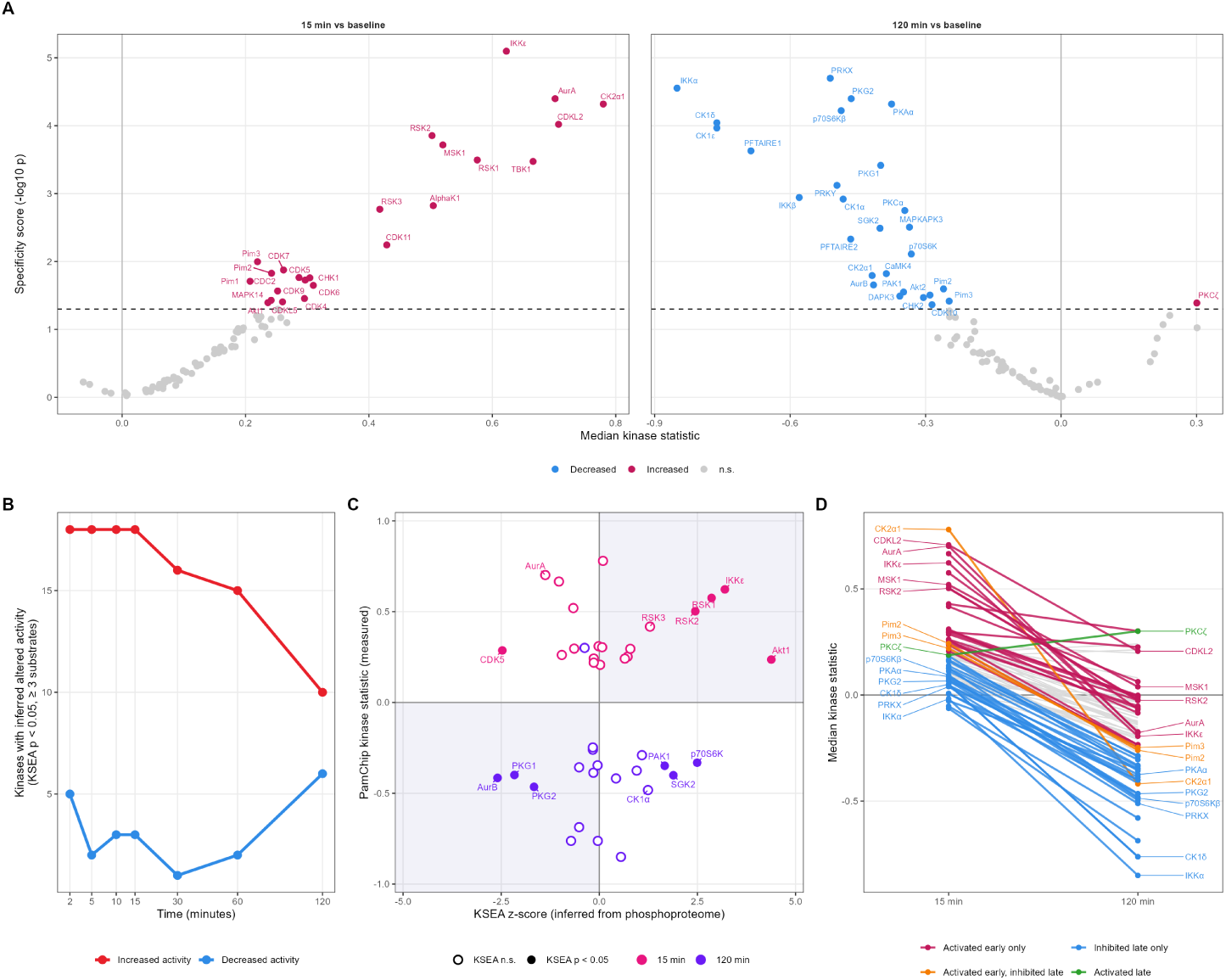
Kinome profiling revealed broad suppression of kinase activity in the late window. (A) Experimental kinase profiling with the PamChip STK peptide array in neutrophils stimulated with *S. aureus* for 15 minutes (left) and 120 minutes (right) versus baseline, n = 10 donors after quality control. Specificity score (*−* log_10_ *p*) against median kinase statistic; red, increased activity; blue, decreased; grey, not significant. The dashed line marks the significance threshold (final score *≥* 1.3). (B) Number of kinases with altered inferred activity by kinase-substrate enrichment analysis of the phosphoproteomic time course, at each timepoint versus baseline (*p <* 0.05 and at least three annotated substrates). Red, increased; blue, decreased. (C) Concordance between inferred and measured kinase activity: KSEA z-score from the phosphoproteome against PamChip kinase statistic, at 15 minutes (red) and 120 minutes (blue). Filled points, KSEA *p <* 0.05; open points, not significant. Shaded quadrants indicate disagreement in direction. (D) Example median kinase trajectories between windows on PamChip STK peptide array

At 15 minutes, enriched kinases effect actin remodelling, oxidase assembly, and cell survival. Enrichment of Rac effector kinases PAK1, PAK2, and PAK3 reflected the pro-phagocytic character of the early window described above. AKT1 was the most strongly supported inferred kinase in the dataset, alongside AKT2, AKT3 and PDK1. RPS6KA1 and RPS6KA3 act downstream of ERK to modulate mTOR signalling and promote cell survival [36] and the upregulation noted is consistent with prominent MAPK1 and MAPK3 phosphorylation described in the early window. The protein kinase C isoforms PRKCB and PRKCD directly induce cytosolic NADPH oxidase component recruitment and assembly [37].

At 120 minutes, KSEA inferred decreased activity of six kinases: CDK5, PRKG1, CAMK2D, AURKB, MAK, and MAP2K7. The best supported are AURKB (*Z* = *−*2.59, 11 annotated substrates), CDK5, (inferred from 14 annotated substrates) and PRKG1 inferred from VASP, the standard readout of cGMP-dependent kinase activity. AURKB and PRKG1 were independently reproduced on the kinome array. Both act on cytoskeletal regulators, CRMP2 on microtubules, VASP on actin, so their downregulation parallels the reduced cytoskeletal activity in the late time window.

In the proteome we noted late decline in FOS and JUNB abundance from their 60-minute peak, on a background of a strong inflammatory output including accumulation of CXCL8 protein (Figure 1G). The decline of the AP-1 subunits at the protein level is not explained by reduced ERK signalling: although its phosphorylation had returned to baseline, pathway output remained inferred as active at 120 minutes. FOS and JUNB are short lived [32, 38] suggesting that the transcription factor is degraded whilst its downstream targets are increasing. A kinase-driven change in the phosphorylation of pre-existing proteins early on was therefore echoed by an increase in the later relevant outputs; meanwhile the pro-inflammatory transcription factor burst decays and driver kinases are withdrawn. The distinct, albeit related, cohorts of kinases enriched in the early window and depleted in the late window suggest that rather than a passive return to baseline via cellular exhaustion, this is a programmed response. These reciprocal changes between early and late windows implicate a model in which programmed kinase deactivation leads to termination of inflammatory pathways once they have been initiated, possibly as an adaptive strategy to limit further activation and mitigate host damage.

### 2.5 Functional Outputs Correlate with Phosphorylation trajectories

Although the temporal clustering described above includes some functional annotation, this was not fine-grained enough to identify potential novel mediators of specific functions during phagosomal formation and maturation [9]. Five neutrophil functions were assayed simultaneously in the neutrophils undergoing the phosphoproteomic time-course analysis: particle engagement (fraction of cells associated with bacterial particles), phagocytic load (quantity of bacterial particles ingested), phagosomal acidification, ROS activity, and elastase activity. All increased monotonically across the timecourse, with variable kinetics and differently timed peak activity (Figure 4A). Engagement saturated early and plateaued, whereas load continued to accumulate more gradually. Thus, cells engaged particles early and then continued to ingest, with a small minority of the population never engaging at all. Phagosomal acidification showed the slowest rate-of-change whilst the kinetics of elastase and ROS activity appeared bimodal. The kinetics of ROS activity may reflect an initial peak of activity from membrane-localised pre-assembled NOX2, with the second smaller peak possibly capturing ongoing phagosomal NOX2 assembly and activity titrated to demand [39, 40].

**Fig. 4.**
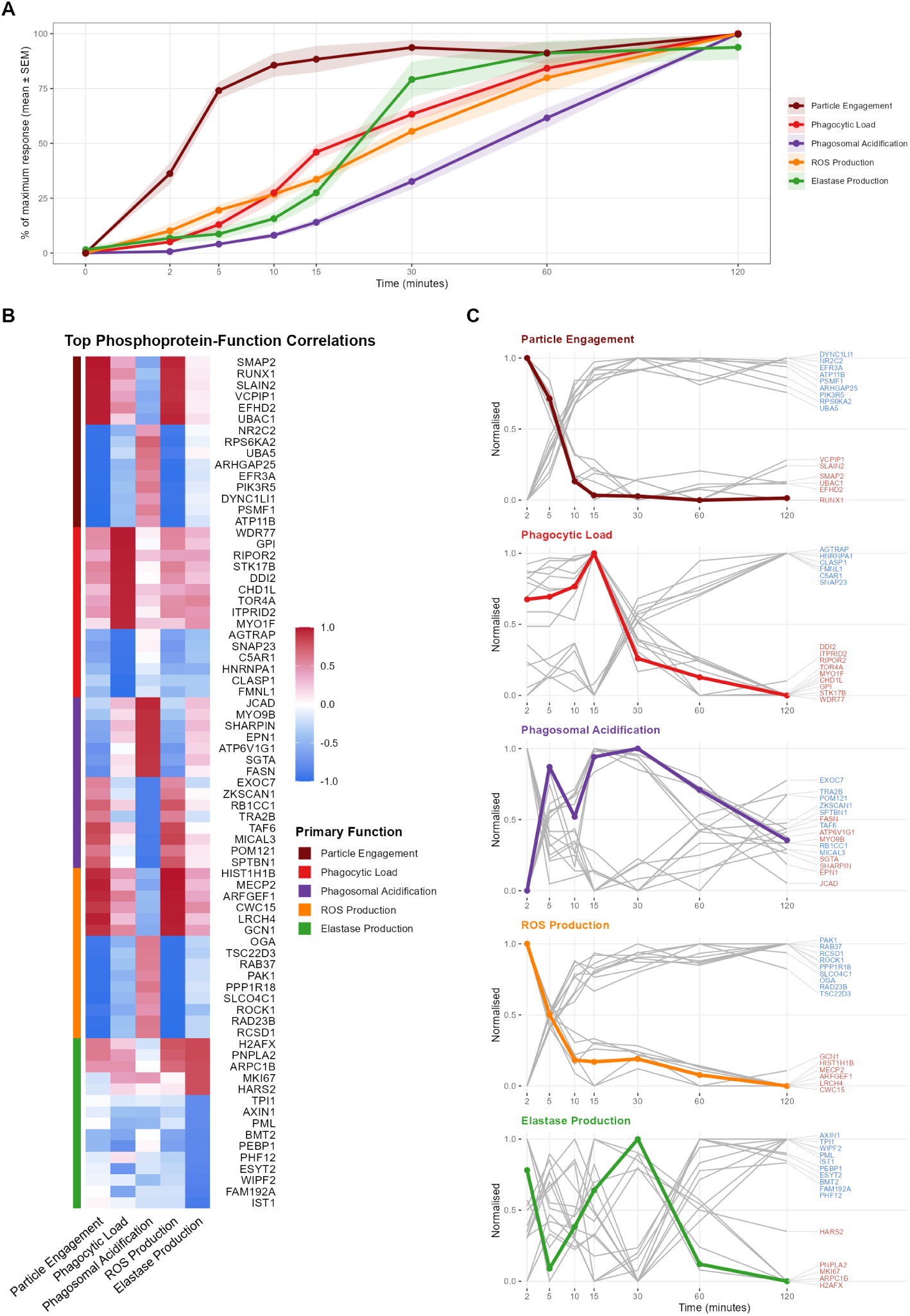
Functional outputs correlate with phosphorylation trajectories (KINETIC). (A) Five neutrophil functions assayed simultaneously in the cells undergoing phosphoproteomic profiling: particle engagement, phagocytic load, phagosomal acidification, ROS production and elastase production, expressed as percentage of maximum response; mean *±* SEM, n = 4 donors. (B) The fifteen phosphoproteins most strongly correlated with each functional velocity curve. Cell colour gives the Pearson correlation between the phosphoprotein trajectory and the velocity curve; the left-hand bar assigns each protein to the function it correlates with most strongly. (C) Normalised trajectories of the correlated phosphoproteins for each function (grey) against the functional velocity curve itself (coloured).

To identify candidate mediators of the five functions, we developed Kinase Network Inference through Trajectory Integration and Correlation (KINETIC) analysis to examine correlations between the trajectories of each phosphoprotein and the functional kinetic curves. Assessed against an empirical null, three of the five functions yielded correlations significantly beyond chance: 125 phosphoproteins reached *r ≥* 0.9 with ROS activity and 113 with particle engagement, against 1̃0 expected by chance; 76 correlated with phagocytic load against 9 expected.

Among the most strongly load-correlated phosphoproteins were the atypical myosin MYO1F, necessary for phagocytic cup formation and closure [41]; CAPG, the actin-capping protein regulating lamellipodial extension [42]; and MSN, the actin–membrane linker essential for cell-shape changes [43]. Overall, 13 of the phosphoproteins identified had known roles in phagocytosis, whilst a further 24 are linked to related processes such as endocytosis and cytoskeletal organisation, however the great majority have no known role and many map to nuclear processes.

The phosphoprotein trajectories most closely tracking particle engagement overlapped almost completely with those correlating with ROS production, reflecting the burst kinetics exhibited by both effector functions: 93% of engagement-correlated phosphoproteins were also ROS-correlated. Those with known neutrophil functions fell into broad categories of activation (including receptor engagement, adhesion and degranulation), phagocytosis (e.g. WASP and its interactor WIPF1 [44]) and ROS production. WASP phosphorylation peaked early and by 30 minutes returned to baseline before becoming progressively dephosphorylated by 120 minutes. Notably there was almost no overlap between engagement and the species in the phagocytic load trajectory, implying distinct mediators of initiation and ongoing ingestion. ROS activity was strongly inversely correlated with the cytosolic NADPH oxidase component NCF2 (p67*^phox^*) and with the haematopoietic Rac-GAP ARHGAP25 [45], consistent with progressive phosphorylation of these proteins as ROS production decelerates in the late window.

Phagosomal acidification was strongly correlated with phosphorylation of the vacuolar proton pump subunit ATP6V1G1, which we have previously demonstrated to be dephosphorylated under conditions of excess exposure to C5a, causing impaired phagosomal acidification [25]. Notably, 40% of the proteins on the acidification list were also present on the ROS list, but all had correlations with inverted signs. The ’antagonistic’ nature of NADPH oxidase activity with acidification, consuming protons, altering v-ATPase membrane insertion and opposing electron and proton flows are well described in neutrophils [46] with inverted correlations indicating a known biological phenomenon.

The recovery of canonical mediators of each function at the top of each correlation list validates this approach, positioning the remaining strongly correlated proteins as putative novel mediators of the respective neutrophil effector functions.

### 2.6 A genome-wide CRISPR knockout screen for phagocytic load recovers the particle engulfment machinery and reveals an antioxidant brake

While KINETIC identifies candidate mediators of function, it cannot confirm requirement. We used <u>C</u>lustered <u>R</u>egularly Interspaced <u>S</u>hort <u>P</u>alindromic <u>R</u>epeats (CRISPR) Cas9 technology to conduct a genome-wide knockout screen in neutrophil-like PLB-985 cells (Figure 5B) sorting on pHrodo fluorescence as a marker of phagocytic load and recovering genes whose loss reduces phagocytosis (phagocytic facilitators, 66 at FDR *<* 0.05) or raises it (phagocytic brakes, 6 at FDR *<* 0.05), FDR thresholds were relaxed for pathway mapping (*<* 0.25 for facilitators, *<* 0.1 for brakes, selecting cut-offs by comparing thresholds for retrieval of canonical phagocytosis mediators and irrelevant, non-immune genes).

**Fig. 5.**
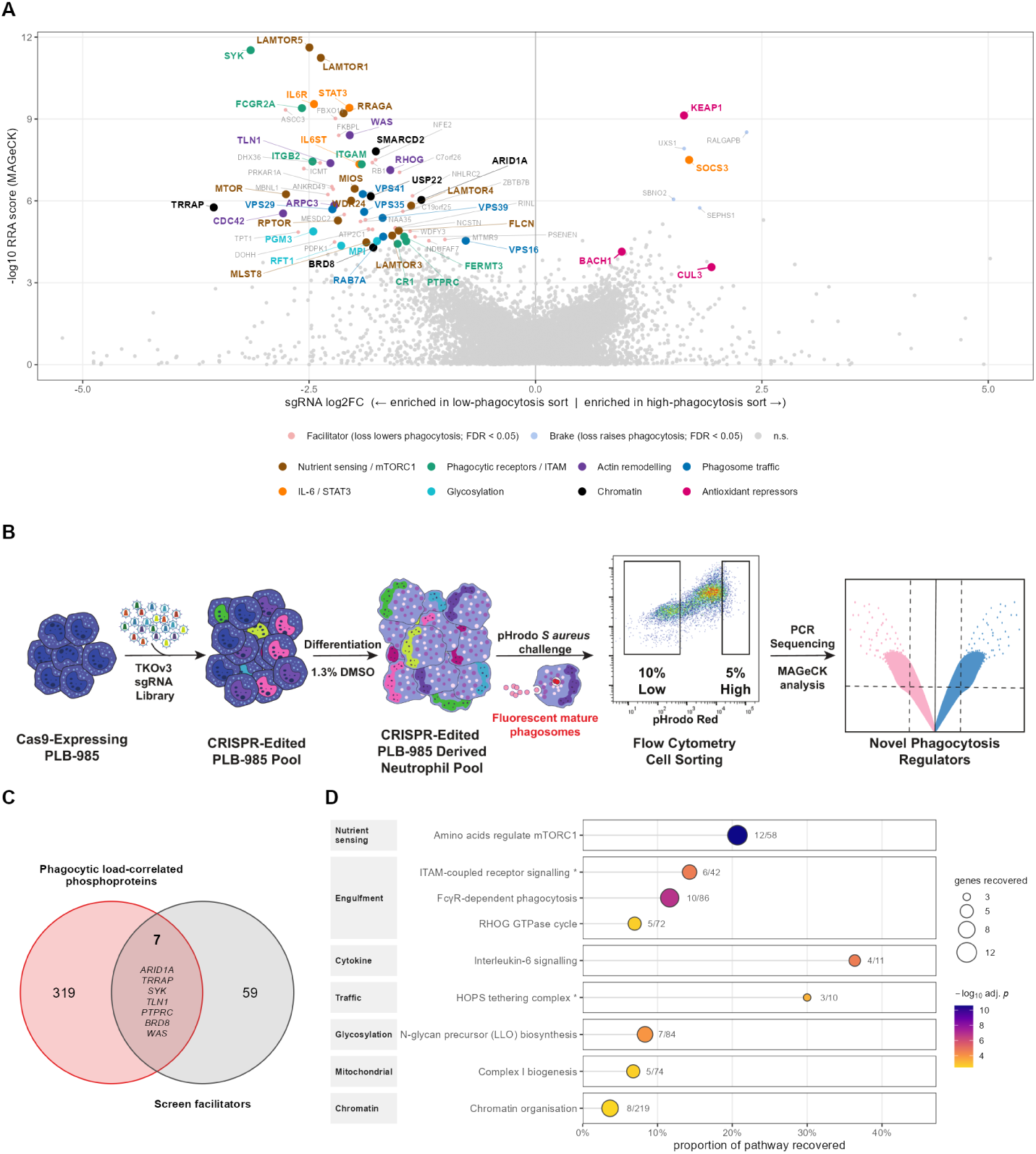
A genome-wide CRISPR knockout screen for phagocytic capacity recovered the engulfment machinery and identifies a redox brake. (A) Genome-wide knockout screen in neutrophil-like PLB-985 cells (TKOv3 library, 18,045 genes), sorted on ingestion of a single *S. aureus* meal. sgRNA log_2_FC between the low- and high-phagocytosis sorts against log_10_(MAGeCK RRA score). Genes whose loss lowers phagocytosis (facilitators, FDR *<* 0.05) and whose loss raises it (brakes, FDR < 0.05) are coloured by pathway module. (B) Screen design. (C) Overlap between the phosphoproteins most strongly correlated with phagocytic load and screen facilitators, with named intersected genes. (D) GO enrichment analysis of CRISPR screen hits, grouped by biological module.

The CRISPR-identified facilitators of phagocytosis included canonical mediators of neutrophil phagocytosis, such as components of the WASP-WAVE-Arp2/3 signalling pathway and the ITAM kinase *SYK*. These corresponded with pathway members identified in the particle engagement (e.g. WASP, WIPF1) and phagocytic load (SYK) KINETIC analysis. Also identified were members of the *β*2-integrin signalling pathway, again demonstrating correspondence between the CRISPR facilitators and KINETIC analysis. Beyond those features identified by both approaches, the screen recovered a broad repertoire of genes coding for canonical neutrophil cell surface proteins including Fc and complement receptors (Figure 5A). The recovery of canonical phagocytosis machinery validated the screen, demonstrating the linkage between the two datasets.

The strongest facilitators identified were the Ragulator components *LAMTOR5* and *LAMTOR1* (also present in particle engagement term in KINETIC), mTORC1 components (*MTOR*, *RPTOR* and *MLST8*), the Rag GTPase *RRAGA* and the regulatory GATOR2 complex (*MIOS* and *WDR24*). Folliculin (*FLCN*), a key GTPase-activating regulator of mTORC1, was also a facilitator and particle engagement KINETIC hit, alongside downstream effectors *EIF4EBP1*, *RPS6*, *RPS6KA2* and *TFEB*. Whilst mTORC2 has well recognised role in actin polymerisation, mTORC1 is less well described in this area, although an mTORC1-RPS6 dependent effect on actin modelling during spermatogenesis has been reported [47].

The other major pathway identified across the CRISPR facilitators and KINETIC analysis was the homotypic fusion and protein sorting (HOPS) complex, including *RAB7A*, which is among both CRISPR facilitators and particle engagement and phagosomal acidification KINETIC hits. HOPS subunits (*VPS16*, *39*, *41*) were well represented in the CRISPR screen alongside retromer components (*VPS29*, *35* and *26C*) and the sorting nexin *SNX17* which plays a role in phagosomal maturation in dendritic cells [48]. The HOPS adaptor *PLEKHM1* was found in the phagocytic load KINETIC, alongside PI3P-responsive *ZFYVE16*. Although the HOPS complex has been implicated in phagosomal-endosome fusion in macrophages, its role in neutrophil phagosomal maturation, where granule fusion is the dominant feature, is poorly understood. Interactions between mTORC1 and HOPS are well described, and these pathways are likely to closely interact in neutrophils as well [49].

The screen also identified the IL-6/JAK/STAT3 pathway: *IL6R*, *STAT3*, *IL6ST* and *JAK1* were all phagocytic facilitators; conversely *SOCS3* which inhibits JAK/STAT signalling downstream of IL6ST was the fourth ranked brake. The internal coherence between screen arms is strong evidence of a role for IL-6 signalling in modulating phagocytosis. This axis was not well represented in the KINETIC analysis or the wider phosphoproteome, and this may be a developmental licencer of phagocytic machinery or a finding specific to PLB-985 cells, although we cannot rule out a direct paracrine effect on mature cells.

Notably, the strongest CRISPR brakes implicated oxidative stress. The top-ranked brake, genome-wide, was *KEAP1* (FDR 0.0012, log_2_FC +1.64, 4/4 guides consistent), the *NFE2L2* (*NRF2*) repressor. Its loss de-represses the antioxidant programme, lowering oxidative stress, which we see here resulted in increased phagocytosis. *BACH1*, a second antioxidant repressor, ranked as 17^th^ brake but did not reach nominal FDR significance. This screen therefore identified the reduction of oxidative stress as a powerful means of increasing phagocytosis. NFE2L2 (NRF2), the master antioxidant transcription factor target of KEAP1, was identified in the phagocytic load KINETIC, and became progressively phosphorylated over the 2-hour time course.

The statistical enrichment of CRISPR screen hits and function-correlated phosphoproteins identified by KINETIC was strongest for the functions related to phagocytosis. Comparing the screen ranks of the 50 most strongly correlated phosphoproteins against the background screen library using a one-sided Mann-Whitney test on facilitator ranks gave *p* = 0.0065 for phagocytic load and *p* = 0.0023 for phagosomal acidification, and 0.088, 0.14, and 0.44 for particle engagement, elastase activity, and ROS activity, respectively. The quantified phosphoproteins ranked higher than average in the screen (median rank 8,101 versus 9,024) so we repeated the comparison against the 1,411 phosphoproteins quantified in both datasets. This attenuated *p* values to 0.065 for phagocytic load and 0.0093 for phagosomal acidification; the remaining three functions stayed non-significant. Against this more conservative reference set the enrichment of load-correlated phosphoproteins was consistent when varying the number of top function-correlated phosphoproteins between 25 and 200, ranging from 0.068 to 0.006. Acidification was less robust. The selective and robust enrichment of load-correlated phosphoproteins among screen hits is consistent with a screen sorting on ingested load, and intersecting the two datasets gives a cohort of high priority candidates.

### 2.7 Transcriptional response of neutrophils to *Staphylococcus aureus*

The phosphoprotein timecourse (Figure 1), notably clusters 8-10 (Figure 2), indicated altered transcriptional activity in *S. aureus*-stimulated neutrophils. We sought to understand how the pathways identified by KINETIC and the CRISPR screens were reflected in the transcriptional response of neutrophils in the early and late time windows (Figure 6). At 30 minutes, 1007 species were significantly altered versus baseline, whereas by 120 minutes this rose to 4188, with 85 percent of genes altered at 30 minutes also altered at 120 minutes. The transcriptional response to stimulation is still expanding while phagocytosis decelerates.

**Fig. 6.**
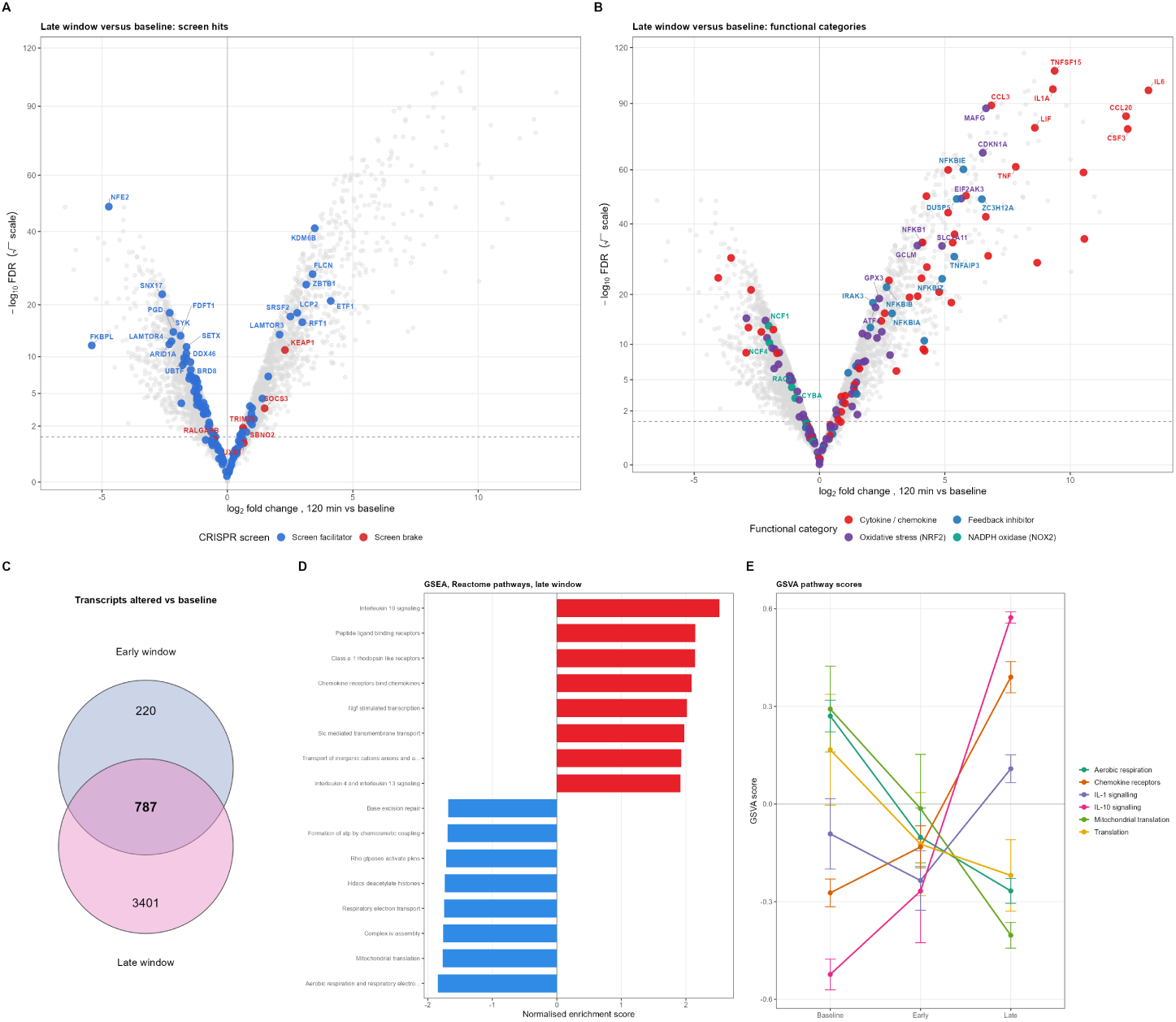
The transcriptional response to *S. aureus* expands into the late window, engaging the antioxidant and inflammatory programmes while downregulating phagocytic machinery. Neutrophils from four healthy donors were stimulated with 50 *µ*g/mL heat-killed, serum-opsonised *S. aureus* bioparticles and sampled in the early (30 minutes) and late (120 minutes) windows. All contrasts were drawn against the pre-stimulation baseline and therefore include the effect of time in culture; the time-matched unstimulated arm was used wherever the stimulation effect itself is quantified in the text. (A, B) Transcripts altered in the late window: log_2_ fold change against *−* log_10_FDR, the latter on a square-root scale. The dashed line marks FDR = 0.05. In (A) points are coloured by CRISPR screen membership (blue, phagocytic facilitator at FDR *<* 0.25; red, phagocytic brake at FDR *<* 0.10; grey, not a screen hit), with all screen hits shown. In (B) the same transcripts are coloured by functional category. (C) Transcripts significantly altered from baseline (FDR *<* 0.05) in each window, with the shared set at the intersection. (D) Gene set enrichment analysis of Reactome pathways in the late window, showing the eight most positively and the eight most negatively enriched pathways by normalised enrichment score; red, positive; blue, negative. (E) Gene set variation analysis scores for six Reactome pathways at baseline and in the early and late windows; mean *±* SEM across four donors.

Notably, all four of the phagocytic brakes identified by the CRISPR screen that were detected in the neutrophil transcriptome and were significantly increased at 120 minutes (*KEAP1*, *SOCS3*, *TRIM28* and *SBNO2*), with *KEAP1* the most induced. Consistent with the signal of a reduction in phagocytic activity in the late window, of the more abundant CRISPR-identified phagocytic facilitators significantly altered in the transcriptome, 69% were repressed. Highly repressed facilitators included the canonical tyrosine kinase *SYK* and novel interactors *SNX17* and *LAMTOR4*.

However, the transcriptome did not point to an entirely anergic phenotype. Consistent with abundance of AP-1 transcription factors in the proteome (Figure 1G), multiple cytokine family signals were increased at 120 minutes including multiple interleukins (e.g *IL-1A* and *IL-1B*, *IL-6*, *IL23A*), downstream signallers from IL1 (e.g. *IRAK2*, *IRAK3*, *NFKB1*, *RIPK2*), chemokines (e.g. *CXCL2*, *CXCL3*, *CXCL8*, *CCL2*, *CCL3*) and TNF family members (e.g. *TNF*, *TNFSF8*, *TNFSF15*) and their inducible proteins (*TNFAIP2,3,6,8*).

The IL-6 pathway identified above to facilitate phagocytosis featured heavily in the late window transcriptome. *IL-6* underwent the strongest transcriptomic induction in the dataset, and LIF and OSM were also induced, as were the receptor subunits *IL6ST* and *IL6R*. *JAK1* and *JAK2* both fell, and counter-regulator *SOCS3* was induced at both timepoints. *ADAM17*, known to shed membrane *IL6R* to release soluble receptor [50], rose four-fold between early and late windows. Together, this suggests that neutrophils produce IL-6 signals while simultaneously reducing their responsiveness to it.

The antioxidant response was induced in the late window, with *NFE2L2*, *MAFG*, *SLC7A11*, *GCLM*, *SQSTM1*, *SOD2*, and *TXNRD1* rising. *KEAP1*, the inhibitor of *NFE2L2* and strong phagocytic brake, rose in concert, consistent with negative feedback of this module. The NADPH oxidase itself is transcriptionally repressed: *NCF1* (p47*^phox^*), *NCF4,*(p40*^phox^*) *RAC2*, and *CYBA* (p22*^phox^*) all fall significantly. Simultaneously, neutrophils exhibited an increased glycolytic signature, with *HIF1A*, *SLC2A1*, *PFKFB3*, *HK2*, and *LDHA* induced, and reduced oxidative phosphorylation signature, suggesting ROS is generated predominantly via NOX2.

The transcriptome therefore supports our contention of a phenotypic shift, with prolonged pathogen exposure resulting in the neutrophil moving from an active phagocyte to pro-inflammatory signaller, activating and recruiting nearby cells.

### 2.8 Integrating phosphoproteomic, KINETIC, transcriptomic, and CRISPR layers identifies key pathways for interrogation

The layers of the *S. aureus* response described above reflect distinct phases of cellular activity. The phosphoproteome captures post-translational modifications of pre-existing proteins. The dynamics capturing changes closer to real time as the phenotypic shift is occurring and KINETIC reports temporal correspondence between groups of phosphoproteins and cells’ functional output. The transcriptome elucidates which phenotype the cell is moving toward, while protein abundance changes are slower. Finally, the CRISPR screen identifies which genes, and hence proteins, modulate phagocytic load. Overlap between layers was sparse, largely because the quantified phosphoproteome covered only 8% of the screen library. Nonetheless, convergence between the layers is identifiable at the module level.

The CRISPR screen identified three main modules as positive determinants of phagocytic load that are supressed during the phenotypic shift between early and late windows. Among these, the ITAM-coupled engulfment machinery was the strongest: *WAS*, *SYK*, *FCGR2A*, *ITGB2*, *TLN1*, and *ITGAM* all ranked among the top 20 phagocytic facilitators, reflected clearly by GO enrichment analysis (Fc*γ*R-dependent phagocytosis, adjusted *p* = 1.5*×* 10*^−^*^7^). WAS and SYK were both KINETIC hits and their phosphorylation and transcript abundance fell into the late window. A similarly enriched module was Ragulator-mTORC1, which represents three top facilitators (*LAMTOR5*, *LAMTOR1*, *RRAGA*; GO enrichment adjusted *p* = 2.6*×* 10*^−^*^11^). LAMTOR1 and FLCN tracked engagement and acidification in KINETIC, with the module being remodelled over the timecourse in both the phosphoproteome and transcriptome.

A third module was chromatin remodelling: *SMARCD2*, *USP22*, *ARID1A*, *TRRAP*, and *BRD8* were all facilitators (chromatin organisation GO, adjusted *p* = 0.0036). ARID1A and TRRAP were the two strongest phagocytic load-correlated phosphoproteins in KINETIC and every SWI/SNF and NSL member detected was repressed by the late window. Meanwhile, the NFE2L2 (NRF2) axis contributed the highest-ranked brake in the screen (NFE2L2 inhibitor *KEAP1*) and its effectors were among the strongest signatures in the late transcriptome. This suggests that a cell that has previously phagocytosed suppresses activity of its engulfment machinery to limit further ingestion, as well as supressing its nutrient sensing apparatus and chromatin remodelling machinery, possibly attenuating initiation of transcriptional responses. Importantly, it also suggests oxidative burden is related to phagocytic appetite.

### 2.9 Neutrophils in the late window are refractory to particulate re-challenge

#### Repeat Stimulation impairs phagocytosis and phagosomal acidification in purified neutrophils

The phosphoproteomic timecourse, kinome, and transcriptomic data revealed reduced activity of phagocytic machinery in late window neutrophils. However, this could reflect insufficient phagocytic targets as much as intrinsic cellular deactivation. To test this, we gave neutrophils a second ’meal’, restoring target availability 15 minutes to three hours after the first. The two ’meals’ carried different fluorophore tags, and the second was double-tagged (Alexa Fluor 488/pHrodo red [25]) to distinguish responses to each (Figure 7A). We assayed phagocytosis by measuring particle engagement and phagocytic load; their product reflects overall population-level eating and is described here as ’phagocytic index’. Similarly, we assayed phagosomal acidification, calculating an acidification index by multiplying the proportion of cells acidifying and degree of acidification. These assays were conducted in purified neutrophils, and then in whole blood, comparing the response to a second *S. aureus* meal with that of time-in-culture matched cells given a second ’meal’ only.

**Fig. 7.**
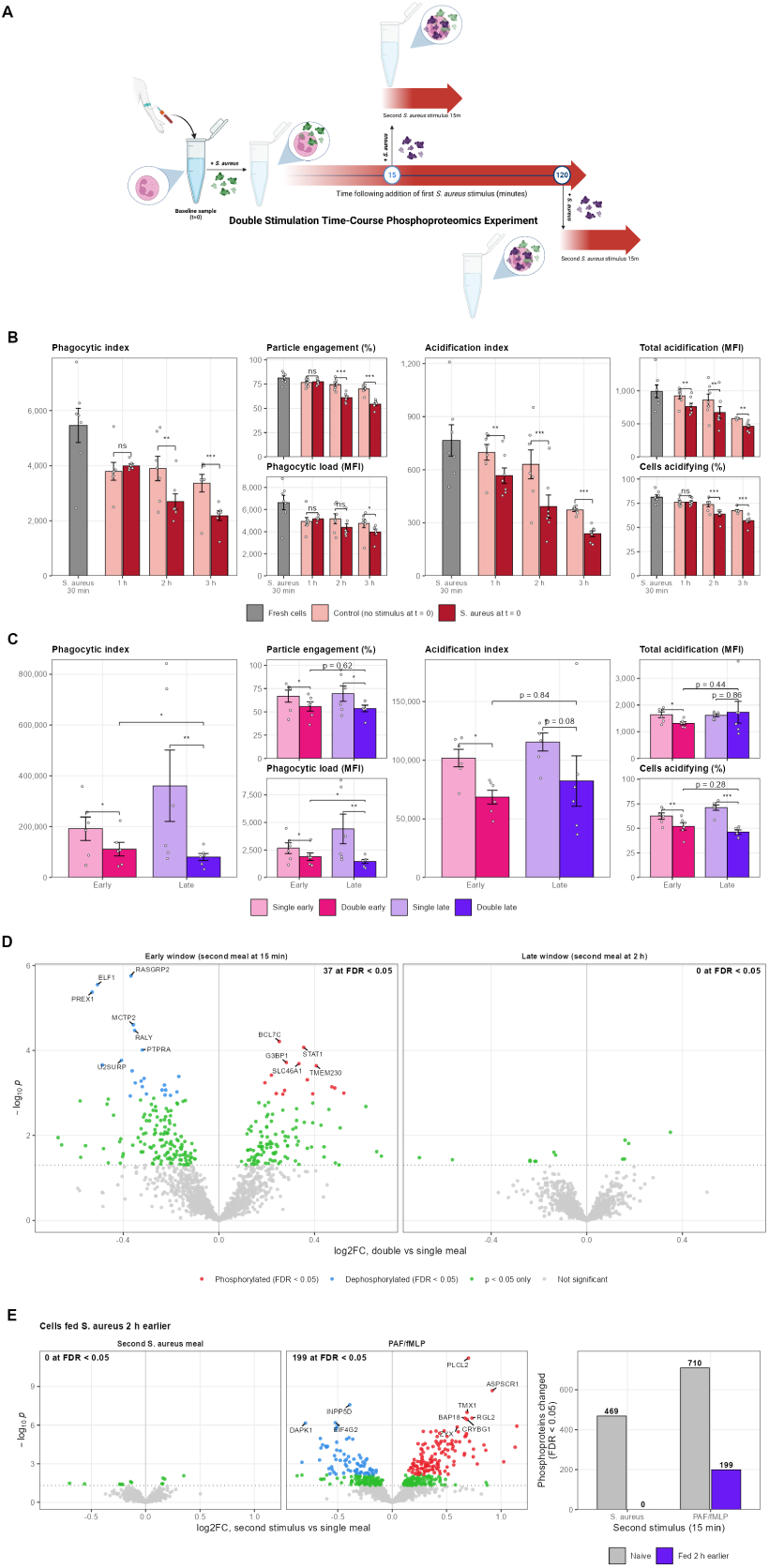
Neutrophils become refractory to repeat stimulation in the late window. (A) Schematic of the design of the double-stimulation experiment: neutrophils received a first *S. aureus* ’meal’ (50 *µ*g/mL) or PBS vehicle control, followed by a 15-minute second challenge with *S. aureus* 15 *µ*g/mL delivered 15 or 120 minutes later. Parallel nonphagocytic experiments primed with platelet activating factor (PAF) and stimulated with formyl-Met-Leu-Phe (fMLP). (B) Purified neutrophils (*n* = 7 donors): phagocytic index, particle engagement, phagocytic load, acidification index, total acidification and proportion of second-’meal’-containing cells that acidify, in cells given a second ’meal’ 15 minutes, 1, 2, or 3 hours after a first, against cells kept *in vitro* for the same time without stimulus. No unstimulated arm was run at 15 minutes; the comparator at this timepoint is freshly isolated cells given a single ’meal’ (grey bar). (C) The same readouts in whole blood (*n* = 6 donors), comparing singly and doubly stimulated cells in the early and late windows. (D) Phosphoproteomic response to the second ’meal’, expressed as doubly versus singly stimulated cells, in the early (left) and late (right) windows: 37 phosphoproteins differ in the early window and none in the late. (E) Cells exposed to *S. aureus* for two hours respond to PAF/fMLP but not to a second *S. aureus* ’meal’ (left), with the number of phosphoproteins altered in naive and previously fed cells for each stimulus (right). Bars show mean *±* SEM with individual donor values; functional comparisons are paired *t* -tests on log-transformed values within donor. *^∗^p <* 0.05, *^∗∗^p <* 0.01, *^∗∗∗^p <* 0.001; ns, not significant.

In purified neutrophils, uptake of the second ’meal’ was completely preserved at short intervals. Neutrophils fed 15 minutes or 1 hour earlier ingested the second ’meal’ as avidly as controls ingested their first, but already showed reduced phagosomal acidification (Figure 7B). In striking contrast, once cells had been exposed to *S. aureus* for 2 hours or more, both ingestion and phagosomal acidification were markedly reduced (Figure 7B). At two hours, prior exposure suppressed ingestion by 31% (*p* = 0.001), deepening to 35% at three hours (*p* = 3 *×* 10*^−^*^5^). This loss was driven by fewer cells engaging particles (−18% at two hours, *p* = 1 *×* 10*^−^*^5^; -22% at three hours, *p* = 2 *×* 10*^−^*^5^), with a smaller fall in the quantity ingested per engaged cell (−15%, *p* = 0.052; -16%, *p* = 0.012). By contrast, phagosomal acidification was impaired from the earliest interval tested: the acidification index fell by 20% at 15 minutes (*p* = 0.001) compared to freshly fed cells, and by 19% at one hour (*p* = 0.005), deepening to 38% at two hours (*p* = 7 *×* 10*^−^*^5^) and 36% at three (*p* = 6 *×* 10*^−^*^4^).

The proportion of cells acidifying their phagosomes fell in step, from 74% to 64% at two hours (*p* = 5 *×* 10*^−^*^4^) and from 67% to 57% at three (*p* = 5 *×* 10*^−^*^4^) (Figure 7B). In purified neutrophils, deficits in phagosomal acidification were therefore the first consequence of serial stimulation, and reduced phagocytic ’appetite’ followed around an hour later. For the first hour after initial encounter, stimulated neutrophils ingested as much as naive cells but acidified their phagosomes less, possibly a state in which internalised bacteria are suboptimally processed. Following this, cells appeared to modulate their appetite and no longer phagocytose as avidly. The damping of both effector functions exceeded the decline attributable to time *in vitro* alone, confirming that it is phagocytic history rather than elapsed time that drives this phenomenon.

To test whether this impairment also occurs in a more physiological milieu, we repeated the assay in whole blood. As we observed a clear effect at two hours and this corresponds to our characterised late window, we compared the responses to second ’meals’ given 15 minutes and two hours after the first. Whole-blood neutrophils largely reproduced the purified-cell phenotype, but unlike purified cells they already ingested less in the early window. In the early (15 minute) window, a prior challenge modestly reduced both components of uptake: the proportion of cells engaging particles fell by 17% (66.9% vs 55.8%; *p* = 0.047) and the quantity ingested per engaged cell by 29% (2,656 vs 1,886; *p* = 0.012), giving a 42% reduction in the composite phagocytic index (*p* = 0.016) (Figure 7C). Acidification was likewise reduced, with the acidification index falling by 33% (101,767 vs 68,469; *p* = 0.013), the proportion of acidifying cells by 17% (62% vs 52%; *p* = 0.008) and acidification per acidifying cell by 19% (*p* = 0.031) (Figure 7C). By the late window the two readouts had diverged: uptake was reduced more strongly, by 78% in phagocytic index (*p* = 0.009). The proportion of cells engaging particles fell by 23% (*p* = 0.021) while the quantity each ingested fell even more markedly by 67%, (*p* = 0.009) (Figure 7C). With acidification the proportion of acidifying cells fell steeply from 71% to 46% (*p* = 2 *×* 10*^−^*^4^), however acidification per acidifying cell was unchanged (*p* = 0.86) (Figure 7C). Prior exposure therefore removed cells from the acidifying population without impairing the acidification of those that remained, whereas for uptake it also reduced the quantity ingested by cells that continued to engage.

We note that whole blood neutrophils in the early window showed a reduction in phagocytic index, whereas purified cells exhibited no such effect. Variation between donors was also greater in whole blood, where the phagocytic index differed up to threefold more between donors than in purified cells. However, the late window effect was striking and consistent: phagocytic index fell in all 13 donors across both experiments. Two donors exhibited atypical patterns: one purified cell donor ingested around 50% more after a early first ’meal’ and one whole blood donor exhibited the strongest reduction in uptake while acidifying more rather than less. These data do not have a sufficient sample size for true inter-donor analyses, but they suggest that how strongly a prior encounter inhibits subsequent phagocytosis varies significantly between individuals.

#### Late window neutrophils are phosphoproteomically refractory to repeat stimulation

With our findings of a substantial phosphoproteomic response to a single *S. aureus* ’meal’ and the striking functional deficits on re-challenge, we characterised the phosphoproteomic response to re-challenge in the early and late windows. A single meal elicited a substantial phosphoproteomic response in both windows, of relatively consistent magnitude: 563 proteins were differentially phosphorylated at 15 minutes versus baseline, and 541 at 120 minutes versus time-matched unstimulated cells (FDR *<* 0.05, of 1,426 quantified). Naive cells given one meal after 120 minutes *in vitro* exhibited altered phosphorylation of 469 proteins relative to the same time-matched unstimulated control. These three responses shared a core of 242 phosphoproteins; with 327 common to the early and late singly stimulated cells and among shared proteins, 98% move in the same direction.

The contrast between the responses to a single ’meal’ in this independent experiment reproduced the early/late distinction described above. The early-specific set contains receptor-proximal phagocytic mediators including the ITAM-coupled phospholipase PLCG2 and its regulator PLCL2, as well as PKD2, MRTFA, and Lamin B1. The late-specific set reflects inflammatory output and simultaneous restraint signals MK2, FOS, FPR2 and the myeloid hexokinase-3 are all phosphorylated late, the latter echoing the HIF-1*α*-driven glycolytic shift seen transcriptionally. The cell also phosphorylated TNFAIP3, PTPN7, IRAK3, ARHGAP1, ARHGAP25, and the ITIM-bearing inhibitory receptors SIRP*α* and CEACAM1. Concurrently, the engulfment machinery was dephosphorylated, including dynamin-2, SPTAN1, DPYSL2, and EFHD2.

Against this background, a second ’meal’ had a diminishing effect. In the early window, a second ’meal’ conferred further alterations in the phosphoproteome: singly and doubly stimulated cells differed from baseline at 563 and 672 proteins, respectively, with fold changes of the shared cohort correlated at *ρ* = 0.855. This finding is caveated in that the doubly stimulated condition was profiled 30 minutes following the first meal while its singly stimulated counterpart was profiled at 15 minutes, conflating the effects of time *in vivo* with those of stimulation. However, in the late window the singly and doubly stimulated states are almost indistinguishable, 541 and 583 features versus the unstimulated time-matched condition, 486 of them shared, with fold changes correlating at *ρ* = 0.955. Direct comparison of the two doubly-stimulated conditions without referencing to unstimulated controls confirmed this observation: 37 phosphoproteins distinguished doubly from singly stimulated cells in the early window, and none in the late window. Across all 1,426 quantified phosphoproteins, with no significance cut-off applied, the typical phosphoprotein changed by 11% in previously unexposed cells, by 7% in cells re-challenged in the early window and by 3% in the late window compared to 8% produced by 120 minutes *in vitro* with no stimulus.

The proteome was less dynamically responsive in general. At FDR *<* 0.05 across 2,983 quantified proteins, 62 differed between doubly and singly stimulated cells in the late window versus the time-matched control, 54 responded to a first ’meal’ after two hours, 14 to a first meal after 15 minutes, and 8 differed with prior exposure in the early window. The direct double-versus-single comparison in both windows yielded no significantly differentially abundant protein. These findings underline the importance of assessing post-translational changes alongside protein abundance analyses.

#### The late phosphoproteomic refractory state is receptor-specific

To distinguish between a globally unresponsive exhausted state and a programmed and specific response to repeat stimulation, we challenged *S. aureus*-stimulated neutrophils with PAF and fMLP, which signal through G-protein-coupled rather than ITAM-coupled receptors. In naive cells the two stimuli evoked largely the same programme. Of the 469 phosphoproteins altered by *S. aureus* and the 710 altered by PAF/fMLP, 425 were shared (91% of the *S. aureus* response) with 201 commonly phosphorylated and 224 commonly dephosphorylated; fold changes correlated at *ρ* = 0.88 across all features quantified.

A naive neutrophil therefore does not distinguish sharply between these stimuli at the level of the phosphoproteome. Conversely, two hours after a first encounter, a second *S. aureus* meal altered no phosphoprotein of 1,426 relative to singly stimulated cells, whilst PAF/fMLP altered 199. These changes recapitulated the naive programme albeit at a reduced magnitude: 158 of the 199 were common to the naive chemoattractant response, 157 of them moving in the same direction, and fold changes across the two correlated at *ρ* = 0.62. The refractory state is therefore not explained by exhaustion; a singly stimulated neutrophil retained the capacity to mount a large and essentially normal phosphoproteomic response to soluble stimuli but not to a bacterial particle.

#### Inferred kinase activity is suppressed on re-challenge in the late window

To identify kinome changes that may underlie the phosphoproteomic refractoriness to serial stimulation in the late window, we conducted KSEA on the double stimulation contrasts, scoring 188 inferred kinases.

In the early window, re-stimulation left the inferred kinase landscape broadly comparable to that of singly-stimulated cells with 13 increased and 8 reduced at nominal significance. In the late window, by contrast, a second meal imposed a clear suppression, reducing 11 kinases and increasing 4. The strongest suppression was of serine/threonine kinases, casein kinase 1*α*, RAF1, CLK1, ROCK1, TTK and the atypical protein kinases C, while the four increased include the protein kinase A catalytic subunits PRKACB and PRKACG (alongside NEK2 and RPS6KA2). This is notable as PKA*α* was among the most strongly inhibited kinases on the kinome array after two hours of a first encounter (Figure 3). Receptor and Src-family tyrosine kinases were also inferred to be suppressed, the group most canonically relevant to particulate receptor signalling; however, fewer than 2.5% of quantified phosphopeptides carry phosphotyrosine, so substrate support for these kinases is sparse.

Despite this, FGR and LYN had *m* values of 4 and 3, respectively, and *SYK* was the second-ranked facilitator of phagocytosis genome-wide above and strongly load-correlated, indicating a potential mechanistic explanation of the ingestion deficit. The ITAM adaptor FCER1G was itself quantified in the phosphoproteome, and followed the same pattern its downstream kinases are inferred to: its phosphorylation rose across a first encounter (log_2_FC +0.47; FDR 9.5 *×* 10*^−^*^4^) and fell on re-challenge (*−*0.24; *p* = 0.04). Across the 107 kinases scored in both early and late double-stimulated conditions, mean inferred activity shifted from +0.37 when the second ’meal’ followed by 15 minutes to *−*0.29 when it followed by 120 minutes (Wilcoxon paired *p* = 1.4 *×* 10*^−^*^4^). The kinases suppressed on re-challenge overlapped with those inhibited at two hours of a single encounter by only 2/11 (*p* = 0.18), and across all 84 kinases scored in both experiments there was no relationship between how far a kinase declined during a first ’meal’ and the extent to which a second suppressed it (*ρ* = *−*0.02; *p* = 0.88). A separate set of kinases differed between the late and early windows in doubly-stimulated cells (Table 3), tracking time since the first ’meal’ rather than the second. AKT1 was amongst these kinases, and was unchanged by re-challenge itself (*z* = +0.13; *p* = 0.45).

**Table 3.**
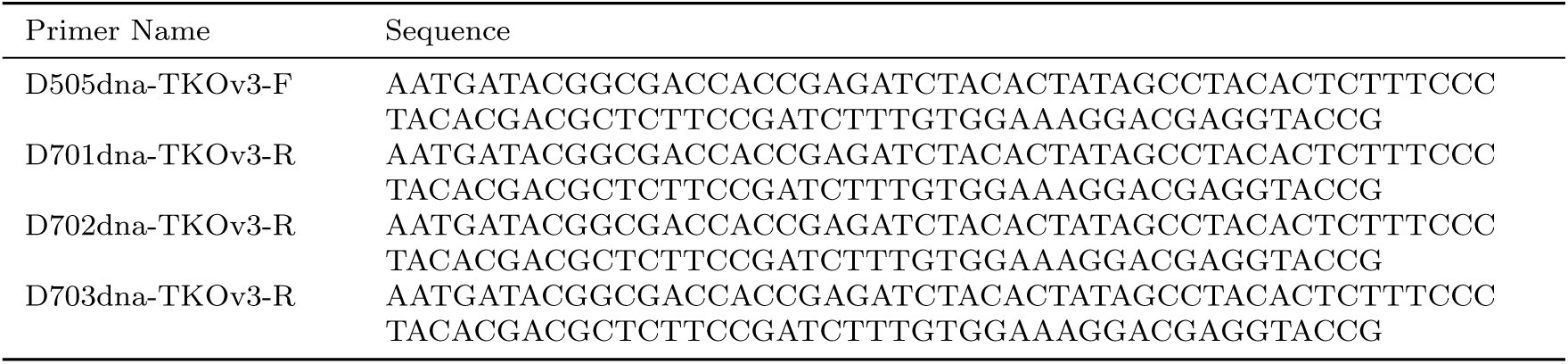
Primers used for inner segment of nested PCR to amplify guide RNA sequences from genomic DNA.

These inferences identify two classes of kinase candidates for future functional interrogation: candidate mediators of the phagocytic rheostat are those kinases whose inferred activity fell after a second ’meal’ and whose function is canonically required for ingestion; restoring them may rescue the phagocytic deficit on re-challenge. Conversely, candidate repressors are those whose activity rose and whose function restrains neutrophil effector function such that inhibiting them may also rescue the deficit.

#### Late window refractoriness spans multi-omic layers

After observing striking phosphoproteomic refractoriness in late window neutrophils, we asked whether this effect operates specifically at the post-translational level or whether it is also echoed across the examined multi-omic layers. We extended the double stimulation experiment to incorporate quantitative proteomics and RNA sequencing across the four conditions.

Firstly, we compared singly and doubly stimulated cells in the late window. In neither ’omic layer did a single feature reached significance: none of 2,980 proteins nor of 13,818 transcripts. With 54 and 5,053 features detected respectively in response to a first ’meal’, this does not appear to be a sensitivity effect. Within these features, a second ’meal’ produced a median effect of 0.06 and 0.20 log_2_ units relative to the first. As with the phosphoproteome, this refractoriness was specific to the late window: in the early window, a second ’meal’ altered 65 transcripts, although no protein, consistent with the slower kinetics of that layer. The phagocytic dose-responsive transcripts in early neutrophils comprised the NF-*κ*B inflammatory programme including *TNF*, *IL1A*, *CXCL2*, *CCL3*, *CCL4*, and *CCL20*, alongside the feedback inhibitors *TNFAIP3*, *NFKBIA*, *NFKBIZ* and *ZC3H12A*. The early window was sampled at 30 minutes rather than the 15 used for the phosphoproteome and the doubly stimulated arm received its second ’meal’ at 45 minutes, so the 65 transcripts responsive to double stimulation in the early window may be inflated due to the extra 15 minutes *in vitro*. This caveat does not apply for the late window: both arms are sampled at 150 minutes. Refractoriness to a second particulate challenge was therefore a property of all three layers, and it developed between the early and late windows.

### 2.10 Features changing between early and late window constitute candidate inputs to the phagocytic rheostat

Comparing singly stimulated and time-matched unstimulated neutrophils in the late window; a single meal alters 534 phosphoproteins, 54 proteins, and 4,110 transcripts beyond the effect two hours *in vitro* alone. Candidate inputs to the rheostat lie in this pool, though membership is necessary rather than sufficient.

The functional data suggest two classes of candidate. Uptake of a second ’meal’ was preserved at one hour and the deficit only arose after two, so could be produced by features that changed significantly between the early and late windows, or by features modulated early and then sustained. Of the pool above, 30,14 and 3,018 features of the above phosphoproteome, proteome and transcriptome pool respecitvely changed between the windows by *≥* 0.5 log_2_FC units in the protein layers or *≥* 1 in the transcript layer. A further 106 phosphoproteins and 773 transcripts were altered early and sustained late.

In the phosphoproteome, switching features included TNFAIP3, ZFP36, PPP1R15A, PTPN7, TAGAP, FOS and the v-ATPase subunit TCIRG1. In the proteome, most began at a lower abundance than baseline and then dwindled further towards the late window; these were granule and secretory proteins including myeloperoxidase, MMP8, lactoferrin, lysozyme, and LCN2, pointing to degranulation over time. FOS and DENND4B alone rose into the late window. Transcripts included *TNFAIP3*, *NFKBIA*, *NFKBIZ*, *ZC3H12A*. These constitute brakes on NF-*κ*B, MAPK signalling, mRNA stability and translation. Furthermore, consistent with an apparent role of oxidative stress, our time-course findings that an NRF2 programme was induced late may be significant.

Given the lag between transcript abundance changes and their manifestation at the protein level, along with the small number of proteins that increase between windows, the changes in the transcriptome are more likely to capture a consequence than an input to the phagocytic rheostat. Of the transcripts, 54 were hits in the CRISPR screen. A stimulated neutrophil therefore continued to receive and transmit inflammatory signals but lost the means to ingest and kill: impaired antimicrobial function with preserved inflammatory output is the combination that characterises neutrophils in sepsis and underlies the paradox of immune dysfunction and hyperinflammation.

### 2.11 Oxidase-derived ROS production constrains appetite of re-challenged neutrophils

The data explored so far supported a role for NADPH oxidase in driving the phagocytic rheostat from multiple angles. Firstly, the oxidase appeared progressively engaged over the interval across which the cell becomes relatively refractory to a second ’meal’ . First, the regulatory subunit NCF2 (p67*^phox^*) was significantly phosphorylated from 10 minutes, remaining so out to 120 minutes, whilst being unchanged following 2 hours of unstimulated *in vitro* incubation. Secondly, redox-sensing machinery was the strongest signal among the phagocytic brakes identified in the CRISPR screen. We therefore hypothesised that ROS accumulated during the first encounter provide the cell with a biochemical record of its phagocytic history, predicting that removing ROS should rescue phagocytosis in doubly but not singly stimulated cells in the late window. To test this, we repeated the whole-blood double-eating assay in the presence or absence of the flavoenzyme inhibitor diphenyleneiodonium (DPI), which abolishes the oxidative burst.

DPI restored phagocytosis in doubly stimulated cells, selectively increasing overall particle ingestion rather than initial engagement in cells re-challenged in the late window, DPI increased the phagocytic index by 124% (*p* = 0.013) (Figure 9A) and total phagocytic load by 95% (*p* = 0.0015) (Figure 9C), while leaving the proportion of cells engaging particles unchanged (+7.9%, *p* = 0.78) (Figure 9B). No such effect was observed in time-matched singly stimulated cells (phagocytic index -16%, *p* = 0.65; load -9.9%, *p* = 0.81). Across all four conditions, DPI saw engagement unaltered (Figure 9B), so a functioning oxidase constrained how much a cell ingests, rather than the decision to engage at all. Importantly, such an effect was not seen in the early window, with DPI having a similar effect on both single and double ’meal’ ingestions.

Phagosomal acidification increased with DPI in every condition (Figure 8A) (median increase in acidification index 3.5- to 4.3-fold), with no dependence on stimulation history. This is consistent with the established role of the oxidative burst in consuming protons and alkalinising the maturing phagosome [46], which is also consistent with the inverted relationship between ROS and phagosomal acidification identified in the KINETIC analysis.

**Fig. 8.**
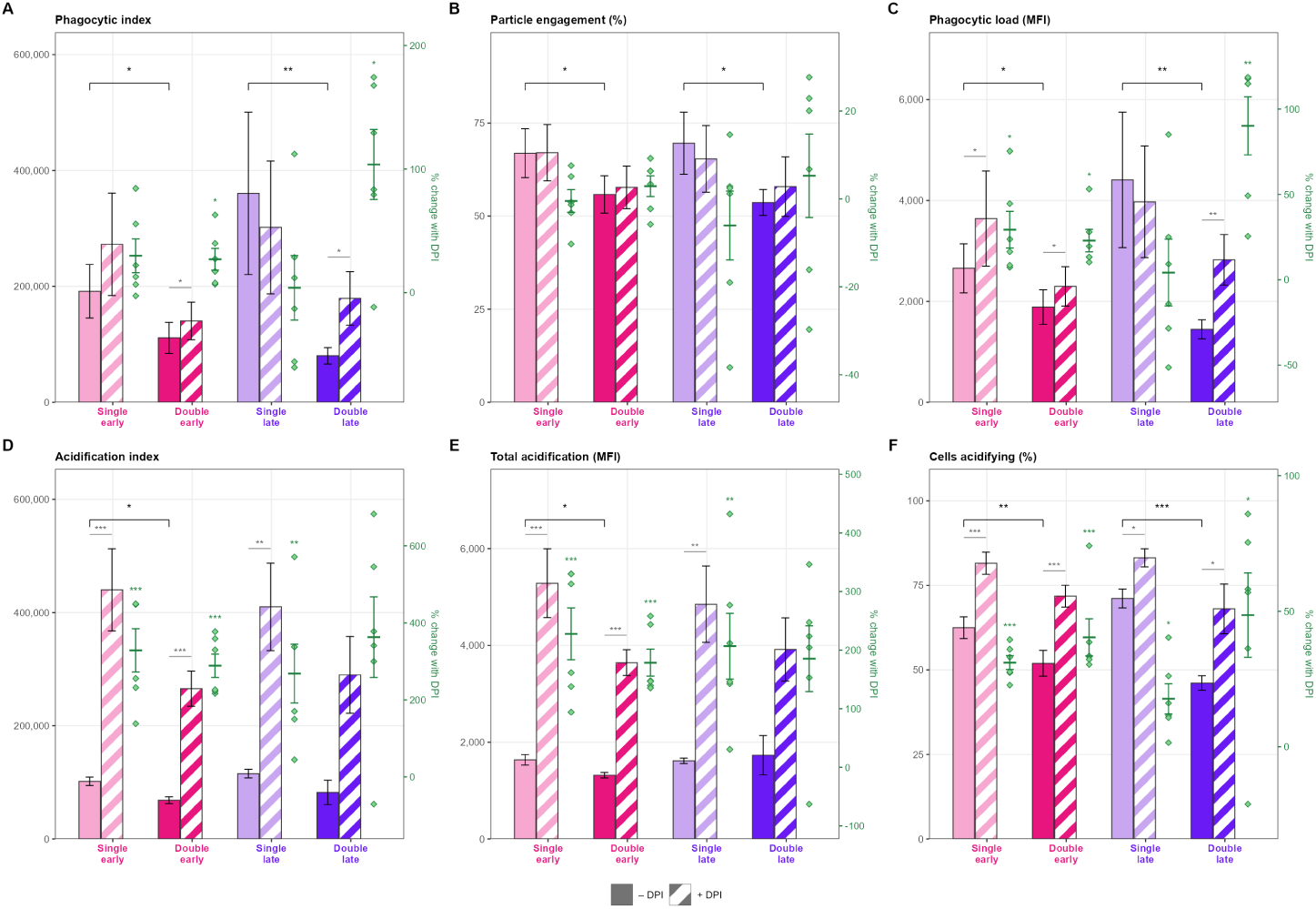
Oxidase-derived ROS constrains the appetite of re-challenged neutrophils. Whole-blood double-stimulation assay repeated in the presence and absence of the flavoenzyme inhibitor diphenyleneiodonium (DPI), n = 6 donors assayed with and without inhibitor. Phagocytic index (A), particle engagement (B), phagocytic load (C), acidification index (D), total acidification (E) and the percentage of cells acidifying (F) are shown for singly and doubly stimulated cells in the early (pink) and late (purple) windows, with (hatched) and without DPI. Bars show mean *±* SEM with individual donor values. Comparisons are paired t-tests on log-transformed values within donor and condition; the dependence of the DPI effect on stimulation history was tested as an interaction on log-transformed per-donor fold changes.

## 3 Discussion

Here, we characterised the neutrophil phosphoproteomic response to a single encounter with *S. aureus*, observing that cells traversed two qualitatively different states: an early window configured for ingestion, in which receptor-proximal and actin machinery was phosphorylated and the kinome broadly activated, and a late window in which that machinery had been supressed, the kinome also suppressed, and the cell reconfigured for inflammatory protein output (Figure 9). Shape-based clustering indicated this to be an ordered progression rather than a switch, with functionally coherent cohorts peaking in sequence from Ras-MAPK signalling at five minutes to degranulation and transcriptional regulation at 120 minutes. The temporal patterns in the phosphoproteome mapped onto those of key effector functions, with highly function-correlated phosphoproteins enriched for canonical mediators of that function alongside putative novel ones. Neutrophils in the late window ingested less, acidified less, and were refractory to re-challenge across the phosphoproteome, proteome, and transcriptome.

**Fig. 9.**
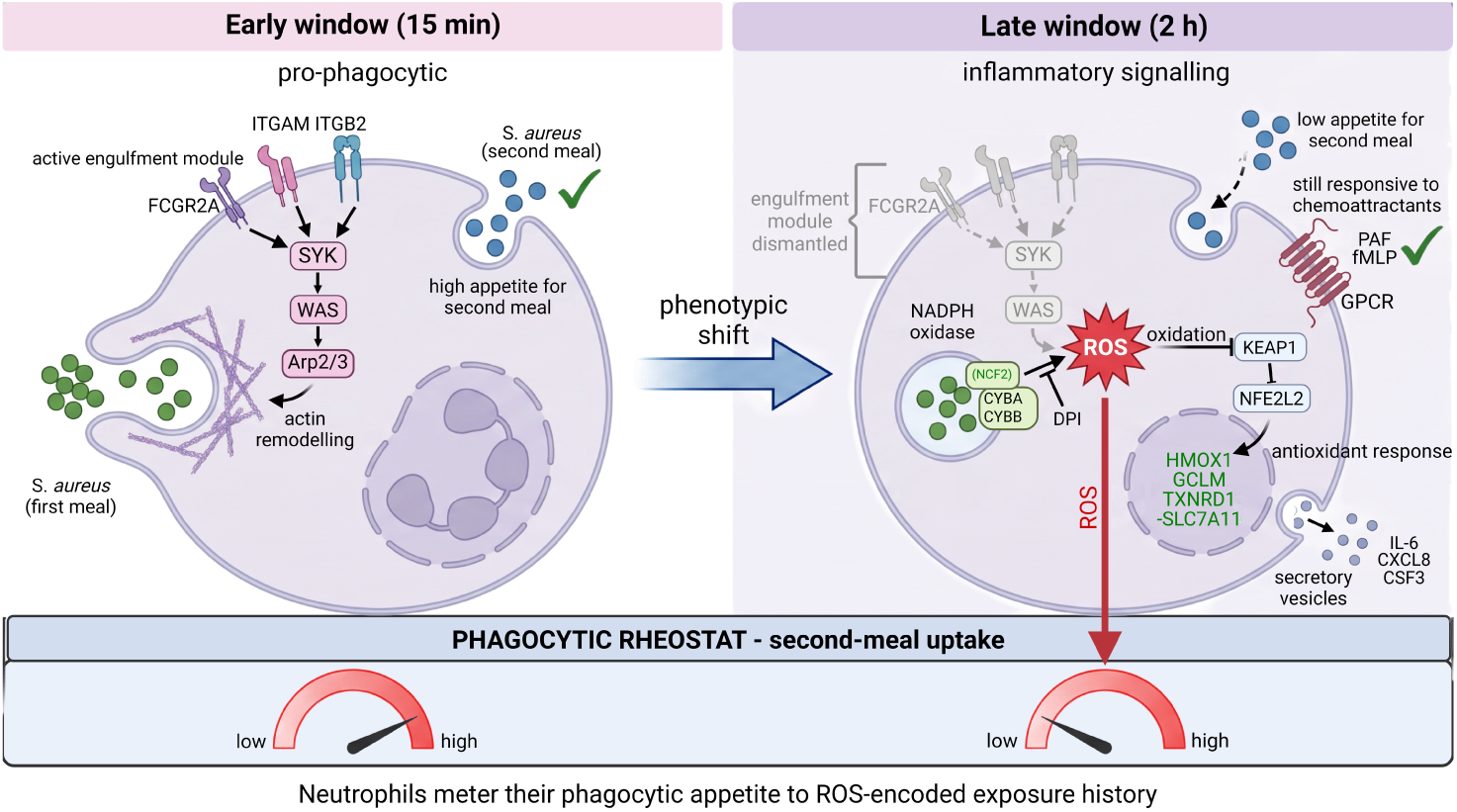
Schematic representation of the interpretation of our data. A ROS-gated rheostat meters neutrophil phagocytosis according to exposure history: schematic representation of putative model. Neutrophils in the early window (left) following a first *S. aureus* ’meal’ (green) retain appetite for a second ’meal’ (blue). By the late window (right) they downregulate their appetite upon re-challenge, on a background of preserved inflammatory output. NADPH oxidase-derived ROS accumulate following pathogen exposure, allowing a biochemical record of phagocytic history and inputting to the phagocytic rheostat. Figure made with Biorender.com

Across all three layers, the late-window cell remained active if relatively refractory to further phagocytosis, and its inflammatory output was amplified: a first ’meal’ induced secreted mediators including *IL-6*, *CSF3*, *CSF2*, *IL-1A*, *TNF* and *CCL20*, each rising more than 30-fold by two hours. Chemokine receptors *CXCR1* and *CXCR2* were strongly repressed, corroborated by less measured at the cell surface on flow cytometry (Supplementary Figure 1). Importantly, we find that the repression is receptor-specific: neutrophils having received a prior *S. aureus* ’meal’ retained their responsiveness to chemoattractant stimuli in the late window (Figure 9). We hypothesise that this phenotype supports recruitment and activation of other leukocytes while the cell downregulates its own capacity to ingest more, possibly in an attempt to reduce the risk of intracellular persistence of viable pathogens.

The timecourse and double stimulation analyses converged on ROS as a modulator of phagocytosis: DPI partially rescued phagocytosis by blocking ROS generation at NOX2, and genetic loss of *KEAP1* raised phagocytosis by de-repressing NFE2L2 and accelerating ROS clearance. The transcriptome showed the antioxidant response was actively engaged in primary neutrophils, and NADPH oxidase subunits transcriptionally repressed. The pharmacological rapid rescue by DPI and the slower timescale of the transcriptional response to stimulation places the ROS-mediated phagocytic deficit predominantly at the post-translational level. The partial nature of the DPI rescue and broad pool of possible rheostat inputs across multi-omic layers above insinuates the rheostat may operate across levels and in a stepwise manner.

Throughout our analyses, the initial engagement of particles and the quantity of particles ingested appeared to be separately regulated. In KINETIC, only 22% of the engagement-correlated phosphoproteins also correlated with phagocytic load. At the functional level, we observed clear divergence too. In purified neutrophils, the late deficit was predominantly caused by fewer cells engaging, whereas in whole blood it was a reduction in phagocytic load per ingesting cell; and DPI rescued load selectively. We therefore propose that the phagocytic rheostat acts to limit appetite. There was notable donor-to-donor variability, especially in the whole blood assay. The mechanisms and consequences of this variability, and their relationship to variability in phagocytic function in patients with severe infection [3, 51] remain to be explored.

This phenotypic plasticity is reminiscent of known forms of innate immune reprogramming. These describe altered responsiveness of innate cells to a second stimulus, but they operate over days, depend on epigenetic remodelling in progenitors, or have been observed cell-intrinsically in longer-lived macrophages and monocytes. Furthermore, they typically manifest as changes in cytokine output. The refractoriness we describe develops within two hours in terminally differentiated neutrophils, constitutes a novel finding with significant translational implications. The CRISPR screen also implicated chromatin remodelling in determining phagocytic appetite with *ARID1A*, *SMARCD2*, *USP22*, and *TRRAP* among facilitators and *ARID1A* the most strongly load-correlated phosphoprotein. *SWI/SNF* members are transcriptionally repressed by the late window.

Impaired phagocytosis on a background of preserved inflammatory output, is consistent with the paradox of sepsis. We have previously described how the anaphlyatoxin C5a acts to impair phagosomal acidification[25], and these data unveil a simultaneous cell-intrinsic route to neutrophil dysfunction. If the phagocytic rheostat we have begun to characterise is an adaptive safeguard to limit uptake of pathogens, in a context of unchecked infection, this could become maladaptive. Notably Hoogendijk and colleagues identified a marked and broad reduction in kinase activity in neutrophils from critically ill patients that is reminiscent of the findings from our study [52].

### 3.1 Limitations

The purified and whole blood neutrophil experiments converged on the late window deficit but diverged early. Differences between the purified and whole blood neutrophil systems may account for this divergence. Whole blood retains opsonins, complement, and other phagocytes that may compete for particles, whereas purification itself can activate neutrophils. The second meal was also given at 15 *µ*g/mL versus 50 *µ*g/mL for the first, so a smaller second response could have a dose effect that we cannot fully exclude here. In the transcriptomic experiment, the early window was 30 minutes rather than 15 minutes, although the kinetics of these layers differ inherently. In transcriptomics and phosphoproteomics experiments, early window singly and doubly stimulated cells were not fully-time matched, as we aimed to give time for the transcriptional response to become apparent in the early window. Finally, donor numbers were relatively small, and the findings should be validated in larger cohorts. Our use of heat-killed *S.aureus* enabled multi-function fluorescent reporter labelling and consistent dosing, but does not fully capture the interactions between live eukaryotic and prokaryotic cells. All work has been conducted in healthy donors or cell lines, and whilst this removes the confounders of illness heterogeneity, at present we cannot determine if similar mechanisms are at work in those with severe infection or other forms of acute illness.

### 3.2 Future directions

Future work should prioritise a CRISPR screen that sorts on the phagocytosis of a second meal, as this will reveal candidate inputs to the rheostat. A suppressive signal acting specifically on re-challenge cannot be recovered during an assay with only one meal. We speculate that rheostat inputs are likely to be post-translational, as transcription occurs on a slower timeline than this phenomenon necessitates; blocking transcription and/or translation during initial *S. aureus* exposure would test this. Transferring medium from fed to naive cells would allow us to isolate whether this response is governed cell intrinsically or extrinsically by secreted factors. Increasing ROS production in the early window would allow us to see whether the deficit could be produced precociously. Furthermore, this work nominates candidate kinases for functional manipulation: PRKG1 and AURKB activity falls in the late window captured by both direct measurement and KSEA inference. If their loss contributes to the deficit, inhibition in the early window should bring it forward. Conversely, their aberrant expression in the late window may delay or rescue the deficit. Meanwhile, an opposite set of PRKACB, PRKACG, NEK2 and RPS6KA2 all increase on serial stimulation in the late window; inhibiting them here may rescue the deficit. Furthermore, unbiased methods to profile the oxidised cysteine proteome have been recently developed [53] that enable the unbiased interrogation of the oxidised proteome for candidate brakes warranting further mechanistic investigation.

## 4 Methods

### 4.1 Donor recruitment and ethics

Donors were recruited voluntarily. Inclusion criteria were adults aged 18 or older who could provide informed consent. Exclusion criteria include known anaemia, pregnancy, recent substantial blood donation, or exposure to transmissible infections such as HIV or hepatitis. Phlebotomy was carried out using a 19G butterfly needle from a vein in the antecubital fossa. Ethical approval was from the Human Biology Research Ethics Committee of the University of Cambridge (HBREC2019.15) and the University of Cambridge acted as sponsor.

### 4.2 Neutrophil purification

The process of density centrifugation for obtaining minimally activated neutrophils has been described previously [54]. Briefly, blood was collected into 4% sodium citrate (1% final concentration). Plasma was removed by centrifugation and erythrocytes depleted by dextran sedimentation. The leukocyte-rich layer was aspirated and separated over discontinuous 42%/51% plasma–Percoll gradients. Granulocytes were recovered from the 42%/51% interface, washed, counted, and assessed for viability by trypan blue exclusion before resuspension in RPMI-1640 with 10% autologous serum. Purity was confirmed by Diff-Quik-stained cytospin (*≥*100 cells counted; target *≥*95% neutrophils) and by flow cytometry as the proportion of violet-high side-scatter events positive for CD16.

### 4.3 *S. aureus* bioparticle preparation and dosing

Heat-killed *S. aureus* bioparticles conjugated to Alexa Fluor 488, Alexa Fluor 405 or Alexa Fluor 594 were used as indicated. The first-stimulus dose of 50 *µ*g/mL was selected by titration across 1.6, 10, 50 and 100 *µ*g/mL, assessing percentage of cells positive, particles ingested per cell, acidification and ROS production over time; 50 *µ*g/mL produced a saturating response not exceeded at 100 *µ*g/mL, corresponding to a multiplicity of infection of approximately 1.5. Second stimuli were delivered at 15 *µ*g/mL, a lower dose chosen so that differences between conditions were not masked by chance encounter at saturating particle:cell ratios.

### 4.4 Functional assays

Purified neutrophils (10 *×* 10^6^ cells/mL in RPMI-1640 with 10% autologous serum) were stimulated with probe-conjugated bioparticles and sampled at 2, 5, 10, 15, 30, 60 and 120 minutes, with baseline controls receiving PBS. Samples were placed on ice at each timepoint to arrest activity, stained with anti-CD16-PerCP for 30 minutes on ice, and acquired on an Attune NxT cytometer. Four functional probes were read simultaneously: Alexa Fluor 488 (particle uptake), pHrodo Red (phagosomal acidification), CellROX Deep Red (ROS activity) and a neutrophil activation probe (elastase activity [55]).

Particle engagement was reported as the percentage of neutrophils positive for Alexa Fluor 488-conjugated bioparticles. Phagocytic load was the median fluorescence intensity of the Alexa Fluor 488-positive population with phagocytic index the product of these two measures. Acidification is reported analogously as the percentage of pHrodo-positive cells and the median pHrodo intensity of that population.

Data are shown as individual donor values with summary statistics. Normality was assessed by D’Agostino–Pearson or Shapiro–Wilk and parametric or non-parametric tests applied accordingly. Comparisons of three or more groups used one-way analysis of variances (ANOVA) with Tukey’s multiple comparisons test. Re-challenge and DPI comparisons are paired within donor and performed on log-transformed values, since per-donor signal scales differ by more than an order of magnitude. Significance was set at *p <* 0.05, two-sided throughout.

### 4.5 Whole blood assay

Whole blood from healthy donors was collected into argatroban (1.5 mg/mL final), which preserves complement and neutrophil function[25], and 200 *µ*L aliquots were dispensed for each condition in triplicate. Aliquots were equilibrated at 37*^◦^*C with shaking at 300 rpm for 10 minutes before stimulation; unstained and cold controls were held on ice throughout. The first ’meal’ was heat-killed *S. aureus* bioparticles conjugated to Alexa Fluor 405, delivered to a final concentration of 100 *µ*g/mL. The second meal was bioparticles conjugated to Alexa Fluor 488, or to pHrodo Red for acidification measurements, delivered at 50 *µ*g/mL either 15 or 120 minutes later. The differing concentrations compared to the purified neutrophils reflected a reduction in apparent phagocytic avidity in the whole blood assay. Cells receiving the second ’meal’ alone, at the matched interval, controlled for the effect of time in culture rather than of prior exposure. All tubes were incubated for exactly 15 minutes after the second meal, then transferred to ice and diluted 100-fold in ice-cold PBS to arrest cellular activity.

Samples were acquired on an Attune NxT cytometer as described previously [25]„ with the dichroic and emission filters of the optical block exchanged for acquisition of whole unlysed blood. First-meal uptake was read on VL2 (405nm excitation, 440/50 nm bandpass emission), second-meal uptake and ROS production on BL1 (488 nm excitation, 530/30 nm bandpass emission), and phagosomal acidification on YL1 (561 nm excitation, 585/16 nm bandpass emission). Phagocytic and acidification indices are the geometric mean fluorescence intensity multiplied by the percentage of positive cells, which normalises for cell number and permits comparison between experiments.

### 4.6 Inhibition of ROS production with DPI

The whole-blood double-stimulation assay was repeated in the presence or absence of the flavoenzyme inhibitor diphenyleneiodonium (DPI, 10 uM final), with ROS production reported by dihydrorhodamine 123 (2.5 uM final) read in the BL1 channel (488 nm excitation, 530/30 nm bandpass emission).

### 4.7 Double stimulation experiment

Purified neutrophils were given a first stimulus of 50 *µ*g/mL *S. aureus* bioparticles, or PBS control as the time-matched unstimulated arm, and incubated at 37*^◦^*C for 15, 60, 120 or 180 minutes. A second stimulus was then delivered for 15 minutes: either 15 *µ*g/mL *S. aureus* bioparticles, or PAF (1 *µ*M) for 5 minutes followed by fMLP (100 nM) for 10 minutes. For the *’omics* arms the second stimulus was 15 minutes for proteomics and phosphoproteomics and 30 minutes for RNA sequencing, and the early timepoint in the transcriptomic experiment corresponds to 30 minutes, not 15. Every re-challenge comparison is made against cells of identical history and culture duration that did not receive a second meal.

### 4.8 Quantitative proteomics and phosphoproteomics

Neutrophils (3 *−* 5 *×* 10^6^ per tube) were pelleted at 10,000*×*g for 1 minute at 4*^◦^*C, the supernatant rapidly aspirated, and pellets snap-frozen in liquid nitrogen and stored at -80*^◦^*C. Samples were processed by the Cancer Research UK Cambridge Institute proteomics facility. Cells were lysed in ice-cold 0.5% SDS, 0.1 M TEAB with HALT protease and phosphatase inhibitors, heated to 90*^◦^*C for 5 minutes, sonicated, and clarified at 20,000*×*g. Protein (100 *µ*g per sample) was reduced with 5 mM TCEP, alkylated with 10 mM methyl methanethiosulfonate, and digested with trypsin (1:30 w/w) overnight at room temperature.

Peptides were labelled with tandem mass tags, pooled in equal ratio, and fractionated at basic pH on an XBridge C18 column into 48 fractions. Fifteen per cent of each fraction was reserved for full proteome analysis; the remainder was consolidated into 14 fractions for phosphopeptide enrichment on High-Select Fe-NTA. Full proteome fractions were pooled to 32 and analysed on an Orbitrap Fusion Lumos with MS3-based quantitation (survey 380–1500 m/z at 120,000 FWHM; MS2 CID in the ion trap; MS3 HCD in the Orbitrap at 50,000 FWHM). Phosphopeptide fractions were analysed on a Q-Exactive HF (survey 400–1600 m/z at 60,000 FWHM; MS2 at 30,000 FWHM).

Spectra were searched in Proteome Discoverer 2.1 with SequestHT against human and *S. aureus* UniProt reviewed databases at 1% spectrum-level FDR (Percolator), with TMT and methylthio as static modifications and oxidation and deamidation as variable. Reporter ion signal-to-noise ratios were extracted and protein grouping applied by parsimony.

Peptide and phosphopeptide intensities were median-scaled and aggregated to protein, phosphoprotein and phosphopeptide level. Missing values in the phosphoproteomic data were imputed by *k*-nearest neighbours (qPLEXanalyzer) for features detected in the majority of samples within a plex (*≥*6 of 11) and the majority of plexes (*≥*3 of 4), yielding 1,503 phosphoproteins in the time course.

Plasma-derived proteins were excluded from the proteome on the basis that 123 of 3,460 proteins exceeded log_2_FC > 1 at two minutes — faster than protein synthesis permits — and that these were complement, immunoglobulin, coagulation and acute-phase proteins largely absent from the neutrophil background transcriptome. The remaining 3,337 proteins were referenced to the two-minute timepoint.

Differential analysis used limma linear models on log intensities. For the double-stimulation experiment a single model was fitted across all conditions and contrasts extracted from it, so that every comparison shares one variance estimate; features were required to be quantified in all donors (*n* = 1,426 phosphoproteins, *n* = 2,983 proteins). Significance is Benjamini–Hochberg FDR *<* 0.05 unless stated.

### 4.9 Shape based clustering

Phosphoprotein trajectories were clustered by dynamic time warping shape-based distance using partitional clustering with shape-extraction centroids (dtwclust v2.1.2 in R), separately for the phosphorylated and dephosphorylated arms. The number of clusters was fixed at *k* = 10 (six phosphorylation, four dephosphorylation) on interpretability grounds; consensus clustering over resampled subsets showed the proportion of ambiguous clustering to decrease monotonically with *k* and identified no preferred *k*, so the partition should be read as a summary of a continuum rather than as ten definitive discrete groups. Partition stability was assessed over 100 restarts from randomised initialisations and summarised as mean pairwise adjusted Rand index, with per-cluster stability reported as mean co-classification frequency.

Protein–protein interactions within and between clusters were taken from STRING at medium (*≥*400) and high (*≥*700) confidence. Inter-cluster enrichment was computed against a degree-preserving null generated by edge rewiring, with permutation *p* values and Benjamini–Hochberg correction across the 55 cluster pairs. Community structure within the phosphorylated arm was detected with the Leiden algorithm (igraph, modularity objective) on the STRING-score-weighted induced subgraph. Resolution was swept over *γ* = 0.4–2.0; at each resolution the algorithm was run from 200 randomised seeds and partitions combined by consensus clustering, thresholding the co-classification matrix at *τ* = 0.5 and re-clustering to convergence (AMI *>* 0.999). Gene Ontology biological process over-representation was computed per cluster with the R package clusterProfiler against a whole genome background with Benjamini-Hochberg correction with cluster. Cluster annotations were performed manually informed by a combination of GO enrichment, PPIs, and hub proteins.

Hub proteins were ranked within each cluster by a composite score combining intra-cluster STRING connectivity, magnitude of absolute fold change for that protein across the timecourse, and contribution to the cluster’s GO enrichment via GO alignment score: sum of log_10_q over all significantly enriched GO biological process terms for that cluster containing the protein.

Plasma-derived proteins were retained in the phosphoproteomic dataset since phosphorylation was quantified relative to baseline rather than as abundance. Each protein’s phosphorylation was compared to its own abundance trajectory across the 952 proteins in both datasets. Six proteins showed a carry over signature whereby an increase in both datasets within two minutes and remaining static. These six (KNG1, ALB, APOA2, AHSG, ITIH2, VTN) were excluded from hub ranking and cluster functional annotations.

### KINETIC analysis

For each of the five functional assays a velocity curve was computed as the first difference of the mean signal across donors between consecutive timepoints, divided by the interval in minutes, giving rate of change per minute. Pearson correlation was computed between each velocity curve and the trajectory of each of the 981 phosphoproteins with complete data and non-zero variance.

### Kinase-substrate enrichment analysis

KSEA was performed with the KSEAapp package in R using PhosphoSitePlus and NetworKIN annotation at default thresholds. Kinases are reported with the number of annotated substrates (m) supporting each inference.

### Experimental kinome profiling

Neutrophil lysates prepared in a kinase-activity-preserving buffer were profiled on the PamGene STK PamChip, a porous three-dimensional peptide microarray carrying 144 serine/threonine phosphosites, run on a PamStation platform with fluorescently labelled anti-phospho-serine/threonine antibodies and read at multiple exposures. Twelve donors were profiled at baseline, 15 and 120 minutes; two were excluded after quality control on the basis of atypical baseline kinase activity, which reduced biological coefficient of variation from 29% to 21%. Ninety-two of 142 phosphosites passed quality control. Phosphosite-level differences were tested by one-way ANOVA with Dunnett’s post-hoc test against baseline. Upstream kinase analysis used the proprietary UKA algorithm, matching peptides to human phosphosites by BLAST and querying Reactome, UniProt, PhosphoELM, HPRD, PhosphoSitePlus and PhosphoNET, with significance and specificity scores derived by permutation.

### 4.10 RNA sequencing

Transcriptional profiling was performed on the double-stimulation experiment only; no RNA sequencing of the seven-point time course was undertaken. Neutrophils from four donors were profiled across the twelve conditions (48 samples), each stimulus history paired with its own time-matched control.

Because the transcriptome was sampled in two windows rather than continuously, every prior-exposure contrast is drawn between cells of matched age. The effect of prior exposure in the late window is therefore double staph stim, late against aged naive, single staph stim: both arms are two and a half hours old and both received a meal 30 minutes before sampling, differing only in whether they had eaten before. The fresh single staph stim arm received the same meal without the preceding two hours in culture and is a younger cell, so it is not used for this comparison. Aged naive, single staph stim against unstimulated, late isolates the response of an aged but previously unstimulated cell to a first meal, and double staph stim, late against single staph stim, late isolates the effect of the second meal itself. The equivalent early-window contrast is double staph stim, early against single staph stim, early, and the chemoattractant arms are compared against their own matched controls in the same way.

RNA was stabilised in TRIzol LS and extracted per manufacturer’s instructions, and accepted for library preparation at A260/A230 and A260/A280 *>* 1.8. Library preparation and sequencing were performed by the CRUK Cambridge Institute Genomics Core using an unstranded library preparation kit, generating paired-end reads of 50 bp. Libraries yielded between 22.4 and 34.2 million read pairs per sample, against a target of 15 million.

Read quality was assessed with FastQC v0.12.0. Low-quality 3*^′^* bases and adapter sequence were removed with TrimGalore v0.5.0, discarding reads shorter than 20 bases after trimming; approximately 0.2% of reads were removed per sample. Reads were mapped with STAR v2.7.11b to the Ensembl *Homo sapiens* GRCh38 reference genome using the corresponding Ensembl GTF annotation. Between 74% and 78% of reads mapped uniquely; reads mapping to multiple loci were discarded to avoid false positives in differential expression. Alignment quality metrics were generated with Picard v1.90 and RSeQC. Gene-level counts were generated with HTSeq. Transcript-level quantification was additionally performed with Salmon, but is not used in the analyses reported here.

Differential expression was performed using edgeR version 3.26.5 under R 3.6.1. Genes with low counts were filtered within each comparison, retaining genes with more than 5 counts per million in at least half of the samples tested, or, where group sizes were unequal, in at least as many samples as the smaller group; the number of genes tested therefore varies by contrast, from 5,324 to 6,928. Effective library sizes were computed by trimmed mean of M-values normalisation, and systematic GC content and gene length bias removed with CQN version 1.30.0. Dispersion was estimated per gene alongside a common dispersion, and pairwise comparisons made with a generalised linear model, with donor passed to the model as a blocking factor so that all contrasts are paired within donor. Multiple testing was controlled by the Benjamini–Hochberg method. Principal component analysis and sample heatmaps used DESeq2 version 1.24.1 with regularised log transformation, computed separately within each comparison; these values were not used for hypothesis testing.

Contrasts requiring a shared variance estimate across arms, and all cross-layer comparisons with the proteomic and phosphoproteomic data, were derived from a single linear model (limma) fitted to log_2_ counts per million across all samples, with donor as a blocking factor and filtering applied once across the full sample set rather than per comparison. Transcript thresholds are Benjamini–Hochberg FDR *<* 0.05 with log_2_FC *≥* 1 unless stated; the fold-change criterion is applied to the transcript layer alone, because a substantially larger proportion of the transcriptome than of either protein layer reaches statistical significance.

### Cell culture and differentiation

PLB-985 cells are a promyelocytic cell line capable of differentiation to neutrophils upon exposure to dimethyl sulfoxide (DMSO) [56]. Cells were maintained at 37*^◦^*C and 5% CO_2_ in RPMI-1640 media supplemented with 1 mM sodium pyruvate, 10 mM HEPES, 1 mM GlutaMAX, and 10% foetal bovine serum. Cells were passaged every 48-72 hours, maintaining a density of 2 *×* 10^5^ *−* 1 *×* 10^6^ cells/mL. HEK293T cells were cultured under adherent conditions in DMEM supplemented with 30 mM HEPES, 1 mM sodium pyruvate, 1 mM GlutaMAX, and 10% foetal bovine serum.

### Lentiviral production

PSPAX2 and PMD2.G were a gift from Didier Trono, and the Toronto human knockout pooled library (TKOv3) was a gift from Jason Moffat (Addgene #90294 ; http://n2t.net/addgene:90294 ; RRID:Addgene_90294). 24 h prior to transfection, cells were seeded at 9 *×* 10^4^ cells/cm^2^ in a T 175 flask. On the day of transfection, 26.8*µ*g of PSPAX2, 8.08*µ*g of pMD2.G, and 26.9 *µ*g were complexed with 247.1 *µ*g of linear polyethyleneimine (25 kDa) in 4 mL of Opti-MEM for 10 minutes at room temperature and added to the HEK293T cells. Media were harvested every day for three days, pooled, and concentrated by ultracentrifugation at 25000 rpm for 2 hours at 4°C. The resuspended pellet was frozen at -80°C for long term storage.

### CRISPR knockout screen

PLB-985 cells expressing Cas9 were transduced at a multiplicity of infection of 0.3 by TKOv3 lentivirus by spinoculation at 800g for 2 hours in the presence of 4 *µ*g/mL of polybrene at 32 °C. Cells were rested for 48 hours, selected in 1 *µ*g/mL of puromycin for 72 hours, and then maintained in antibiotic-free complete media. At day 17 post transduction, cells were transitioned to media containing 2% Nutridoma-CS, 1.3% DMSO, and 0.5% foetal bovine serum.

At day 21, pHrodo red-conjugated *S. aureus* bioparticles were opsonised with human serum for 30 minutes at 37°C. These were added at a final concentration of 50 *µ*g/mL to a suspension of 5 *×* 10^6^ differentiated cells/mL and incubated on a rotary shaker for 30 minutes at 37°C. Cells were resuspended at 1 *×* 10^7^ cells/mL on ice and sorted for the top 5% and bottom 10% of phagocytic capacity based on pHrodo-red fluorescence in the 561nm 582/15 channel of the BD FACSAria fuorescent sorter. Using an 85 *µ* nozzle, cells were sorted based on fluorescence of pHrodo red in the 561 nm excitation, 582/15 nm emission into 4 mL polypropylene tubes on ice, and immediately processed for genomic DNA using the Wizard genomic DNA kit according to manufacturers’ instructions. Genomic DNA was amplified by nested PCR using Q5 high fidelity 2X mastermix (New England Biolabs) with outer primer pair GAGGGCCTATTTCCCATGATTC /CAAACCCAGGGCTGCCTTGGAA and inner primer pairs listed in table 3. Input and cycle number for each PCR reaction was titrated empirically and products analysed by agarose gel electrophoresis. PCR products were size selected using SPRIselect beads according to manufacturers’ instructions, and library quality was analysed using the Agilent 2100 Bioanalyzer system with the DNA 1000 kit. The pooled library was sequenced on the Illumina NovaSeq X platform, allocating 50 million reads to parent libraries and 20 million to sorted libraries.

Raw sequencing reads were trimmed to leave the 20 nt guide sequence using Cutadapt and aligned to the TKOv3 library to assign guide and gene identities. Guide RNA enrichment was analysed using Model-based Analysis of Genome-wide CRISPR-Cas9 Knockout (MaGECK) as described previously [57, 58]. This uses the negative binomial distribution using a robust rank aggregation algorithm to calculate gene enrichment or depletion.

Correlation coefficients from KINETIC were compared with gene-level ranks from the CRISPR screen to ask whether phosphoproteins correlated with a given function are enriched among the genes required for phagocytosis. For each function, the MAGeCK ranks from the low phagocytosis sort were retrieved for the 50 top positively correlated phosphoproteins and compared to the ranks of the remaining genes in the library (n=18,046) by one-sided Mann-Whitney test.

## Declarations

- **Funding**: ACM is supported by a Clinician Scientist Fellowship from the Medical Research Council (MR/V006118/1), with work in this manuscript supported by a project grant from the National Institute of Academic Anaesthesia (WKRO-2018-0051). MI is a Gates Scholar, supported by the Gates Cambridge Trust.
- **Conflict of interest/Competing interests**: Not applicable
- **Ethics approval and consent to participate**: Blood from healthy donors was obtained under a protocol approved by the Human Biology Research Ethics Committee of the University of Cambridge (HBREC2019.15).
- **Consent for publication**: Not applicable
- **Data availability**: Phosphoproteomic, proteomic and transcriptomic data from primary human neutrophils are deposited and will be released publicly on acceptance for publication following peer review.
- **Materials availability**: Not applicable
- **Code availability**: Not applicable
- **Author contribution**: Conceptualisation: MRoeder, MI, AYKCN, DCT, KO, ACM Methodology: MRoeder, MI, AYKCN, AJTW, DCT, ACM Formal Analysis: MRoeder, MI, AYKCN, ITW, SL,KK, ACM Investigation: MRoeder, MI, AYKCN, HB, ITW, SL, MH, AS, IM, ML, MRoutledge, KK, CDS, EP, ACM Resources: CDS, DCT, KO, ACM Data curation: MRoeder, MI, AYKCN, HB, MH, CDS, DCT, KO, ACM Visualisation: MRoeder, MI, ACM Writing-original draft: MRoeder, AYKCN, ACM Writing-review and editing: MRoeder, MI, AYKCN, HB, ITW, SL, MH, AS, IM, ML, AJTW, MR, KK, CDS, EP, DCT, KO, ACM Supervision: DCT, KO, ACM Administration: DCT, KO, ACM Funding acquisition: DCT, ACM

## References

[1] Grieshaber-Bouyer, R. et al. The neutrotime transcriptional signature defines a single continuum of neutrophils across biological compartments. Nature Communications 12, 2856 (2021).

[2] Wu, Y. et al. Neutrophil profiling illuminates anti-tumor antigen-presenting potency. Cell 187, 1422–1439.e24 (2024).

[3] Conway Morris, A., et al. C5a-mediated neutrophil dysfunction is RhoA-dependent and predicts infection in critically ill patients. Blood 117, 5178–5188 (2011).

[4] Seo, S. K., Liu, C. & Dadwal, S. S. Infectious disease complications in patients with cancer. Critical Care Clinics 37, 69–84 (2021).

[5] Herrero-Cervera, A., Soehnlein, O. & Kenne, E. Neutrophils in chronic inflammatory diseases. Cellular & Molecular Immunology 19, 177–191 (2022).

[6] Boers, L. S. et al. Alveolar dysregulation of host response in pneumonia and ARDS: implications for immune modulation and infection. Annals of Intensive Care 16, 100085 (2026).

[7] Kwok, A. J. et al. Neutrophils and emergency granulopoiesis drive immune suppression and an extreme response endotype during sepsis. Nature Immunology 24, 767–779 (2023).

[8] Jeffrey, M. et al. Pulmonary inflammation in severe pneumonia is characterised by compartmentalised and mechanistically distinct sub-phenotypes. Nature Communications 17, 5312 (2026).

[9] Naish, E. et al. The formation and function of the neutrophil phagosome. Immunological Reviews 314, 158–180 (2023).

[10] Laskay, T., van Zandbergen, G. & Solbach, W. Neutrophil granulocytes as host cells and transport vehicles for intracellular pathogens: apoptosis as infection-promoting factor. Immunobiology 213, 183–191 (2008).

[11] Gresham, H. D. et al. Survival of *Staphylococcus aureus* inside neutrophils contributes to infection. Journal of Immunology 164, 3713–3722 (2000).

[12] Thwaites, G. E. & Gant, V. Are bloodstream leukocytes Trojan Horses for the metastasis of *Staphylococcus aureus*? Nature Reviews Microbiology 9, 215–222 (2011).

[13] Paclet, M.-H., Laurans, S. & Dupré-Crochet, S. Regulation of neutrophil NADPH oxidase, NOX2: A crucial effector in neutrophil phenotype and function. Front. Cell Dev. Biol. 10, 945749 (2022).

[14] Hakkim, A. et al. Activation of the Raf-MEK-ERK pathway is required for neutrophil extracellular trap formation. Nature Chemical Biology 7, 75–77 (2011).

[15] Azzouz, D., Khan, M. A. & Palaniyar, N. ROS induces NETosis by oxidizing DNA and initiating DNA repair. Cell Death Discovery 7, 113 (2021).

[16] Wang, G. et al. Oxidant sensing by TRPM2 inhibits neutrophil migration and mitigates inflammation. Developmental Cell 38, 453–462 (2016).

[17] Ferling, I., et al. Reactive oxygen species suppress phagocyte surveillance by oxidizing cytoskeletal regulators. eLife (2024). Reviewed Preprint.

[18] Paulsen, C. E., Truong, T. H., Garcia, F. J., Homann, A. et al. Peroxide-dependent sulfenylation of the EGFR catalytic site enhances kinase activity. Nat. Chem. Biol. 8, 57–64 (2011).

[19] Qiu, B., Zhang, S., Ge, S. et al. Vascular smooth muscle cell-derived SO2 sulphenylated interferon regulatory factor 1 to inhibit VSMC senescence. Front. Pharmacol. 16, 1516885 (2025).

[20] Netea, M. G., Domínguez-Andrés, J., Barreiro, L. B. et al. Defining trained immunity and its role in health and disease. Nature Reviews Immunology 20, 375–388 (2020).

[21] Foster, S. L., Hargreaves, D. C. & Medzhitov, R. Gene-specific control of inflammation by TLR-induced chromatin modifications. Nature 447, 972–978 (2007).

[22] Cheng, S. C., Quintin, J., Cramer, R. A. et al. mTOR- and HIF-1*α*-mediated aerobic glycolysis as metabolic basis for trained immunity. Science 345, 1250684 (2014).

[23] Kalafati, L., Kourtzelis, I., Schulte-Schrepping, J. et al. Innate immune training of granulopoiesis promotes anti-tumor activity. Cell 183, 771–785.e12 (2020).

[24] Moorlag, S. J. C. F. M., Rodriguez-Rosales, Y. A., Gillard, J. et al. BCG vaccination induces long-term functional reprogramming of human neutrophils. Cell Reports 33, 108387 (2020).

[25] Wood, A. J. T., et al. C5a impairs phagosomal maturation in the neutrophil through phosphoproteomic remodeling. JCI Insight 5, e137029 (2020).

[26] Zhang, F., Xia, Y., Su, J. et al. Neutrophil diversity and function in health and disease. Signal Transduction and Targeted Therapy 9, 343 (2024).

[27] Buxboim, A., Swift, J., Irianto, J., Spinler, K. R. et al. Matrix elasticity regulates lamin-A,C phosphorylation and turnover with feedback to actomyosin. Current Biology 24, 1909–1917 (2014).

[28] Strzelecka-Kiliszek, A., Kwiatkowska, K. & Sobota, A. Lyn and Syk kinases are sequentially engaged in phagocytosis mediated by Fc*γ*R. Journal of Immunology 169, 6787–6794 (2002).

[29] Majeed, M., Caveggion, E., Lowell, C. A. & Berton, G. Role of Src kinases and Syk in Fc*γ* receptor-mediated phagocytosis and phagosome-lysosome fusion. Journal of Leukocyte Biology 70, 801–811 (2001).

[30] Nardone, C. et al. A heterotrimeric protein complex assembles the metazoan V-ATPase upon dissipation of proton gradients. Nature Structural & Molecular Biology 32, 2076–2087 (2025).

[31] Makita, S., Takatori, H. & Nakajima, H. Post-transcriptional regulation of immune responses and inflammatory diseases by RNA-binding ZFP36 family proteins. Frontiers in Immunology 12, 711633 (2021).

[32] Shaulian, E. & Karin, M. AP-1 as a regulator of cell life and death. Nature Cell Biology 4, E131–E136 (2002).

[33] Miralda, I., Uriarte, S. M. & McLeish, K. R. Multiple phenotypic changes define neutrophil priming. Frontiers in Cellular and Infection Microbiology 7, 217 (2017).

[34] Sardá-Espinosa, A. Time-series clustering in R using the dtwclust package. The R Journal 11, 22–43 (2019).

[35] Paparrizos, J. & Gravano, L. Fast and accurate time-series clustering. ACM Transactions on Database Systems 42, 1–49 (2017).

[36] Anjum, R. & Blenis, J. The RSK family of kinases: emerging roles in cellular signalling. Nature Reviews Molecular Cell Biology 9, 747–758 (2008).

[37] Fontayne, A., Dang, P. M., Gougerot-Pocidalo, M. A. & El-Benna, J. Phosphorylation of p47phox sites by PKC *α*, *β*ii, *δ*, and *ζ*: effect on binding to p22phox and on NADPH oxidase activation. Biochemistry 41, 7743–7750 (2002).

[38] Acquaviva, C., Ferrara, P., Bossis, G., Brockly, F. et al. Degradation of cellular and viral Fos proteins. Biochimie 83, 357–362 (2001).

[39] Li, X. J., Marchal, C. C., Stull, N. D. et al. p47phox Phox homology domain regulates plasma membrane but not phagosome neutrophil NADPH oxidase activation. Journal of Biological Chemistry 285, 35169–35179 (2010).

[40] Quinn, M. T. & Gauss, K. A. Structure and regulation of the neutrophil respiratory burst oxidase: comparison with nonphagocyte oxidases. Journal of Leukocyte Biology 76, 760–781 (2004).

[41] Barger, S. R., Reilly, N. S., Shutova, M. S. et al. Membrane-cytoskeletal crosstalk mediated by myosin-I regulates adhesion turnover during phagocytosis. Nature Communications 10, 1249 (2019).

[42] Witke, W., Li, W., Kwiatkowski, D. J. et al. Comparisons of CapG and gelsolin-null macrophages: demonstration of a unique role for CapG in receptor-mediated ruffling, phagocytosis, and vesicle rocketing. Journal of Cell Biology 154, 775–784 (2001).

[43] Huang, L., Wong, T. Y., Lin, R. C. et al. Replacement of threonine 558, a critical site of phosphorylation of moesin in vivo, with aspartate activates F-actin binding of moesin. Journal of Biological Chemistry 274, 12803–12810 (1999).

[44] Tsuboi, S. & Meerloo, J. Wiskott-Aldrich syndrome protein is a key regulator of the phagocytic cup formation in macrophages. Journal of Biological Chemistry 282, 34194–34203 (2007).

[45] Lőrincz, Á. M., Szarvas, G., Smith, S. M. et al. Role of Rac GTPase activating proteins in regulation of NADPH oxidase in human neutrophils. Free Radical Biology and Medicine 68, 65–71 (2014).

[46] Jankowski, A., Scott, C. C. & Grinstein, S. Determinants of the phagosomal pH in neutrophils. Journal of Biological Chemistry 277, 6059–6066 (2002).

[47] Mok, K. W., Mruk, D. D., Silvestrini, B. et al. rpS6 regulates blood-testis barrier dynamics by affecting F-actin organization and protein recruitment. Endocrinology 153, 5036–5048 (2012).

[48] Dinamarca, S., Croce, C., Salvioni, A. et al. SNX17 regulates antigen internalisation and phagosomal maturation by dendritic cells. Immunology 174, 167–185 (2025).

[49] Hesketh, G. G., Papazotos, F., Pawling, J. et al. The GATOR-Rag GTPase pathway inhibits mTORC1 activation by lysosome-derived amino acids. Science 370, 351–356 (2020).

[50] Rose-John, S. IL-6 trans-signaling via the soluble IL-6 receptor: importance for the pro-inflammatory activities of IL-6. International Journal of Biological Sciences 8, 1237–1247 (2012).

[51] Conway Morris, A., et al. Cell-surface signatures of immune dysfunction risk-stratify critically ill patients: INFECT study. Intensive Care Medicine 44, 627–635 (2018).

[52] Hoogendijk, A. J. et al. Kinase activity is impaired in neutrophils of sepsis patients. Haematologica 104, e233–e235 (2019).

[53] Xiao, H., Jedrychowski, M. P., Schweppe, D. K., Huttlin, E. L. et al. A quantitative tissue-specific landscape of protein redox regulation during aging. Cell 180, 968–983.e24 (2020).

[54] Haslett, C., Guthrie, L. A., Kopaniak, M. M., Johnston, R. B., Jr. & Henson, P. M. Modulation of multiple neutrophil functions by preparative methods or trace concentrations of bacterial lipopolysaccharide. The American Journal of Pathology 119, 101–110 (1985).

[55] Craven, T. H. et al. Activated neutrophil fluorescent imaging technique for human lungs. Scientific Reports 11, 976 (2021).

[56] Rincón, E., Rocha-Gregg, B. L. & Collins, S. R. A map of gene expression in neutrophil-like cell lines. BMC Genomics 19, 573 (2018).

[57] Wit, N., Gogola, E., West, J. A. et al. A histone deacetylase 3 and mitochondrial complex I axis regulates toxic formaldehyde production. Sci. Adv. 9, eadg2235 (2023).

[58] Li, W., Xu, H., Xiao, T. et al. MAGeCK enables robust identification of essential genes from genome-scale CRISPR/Cas9 knockout screens. Genome Biol. 15, 554 (2014).

