## Supplementary figures and images for "Temporal phosphoproteomic programmes set a redox-gated rheostat on neutrophil phagocytosis"

### Supplementary Figure 1

**A**

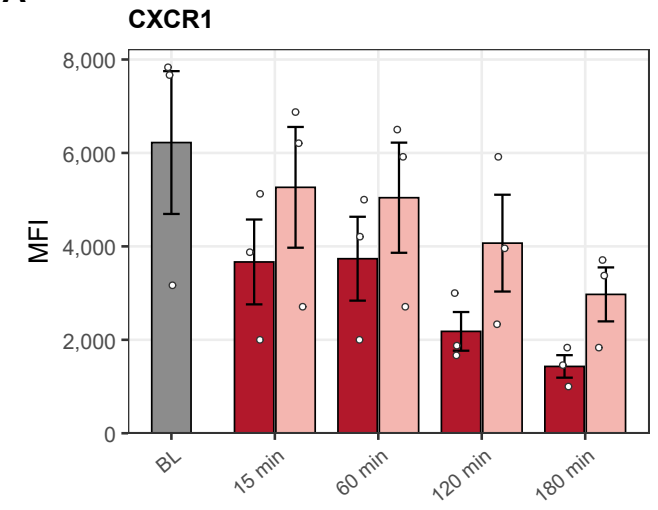

**B**

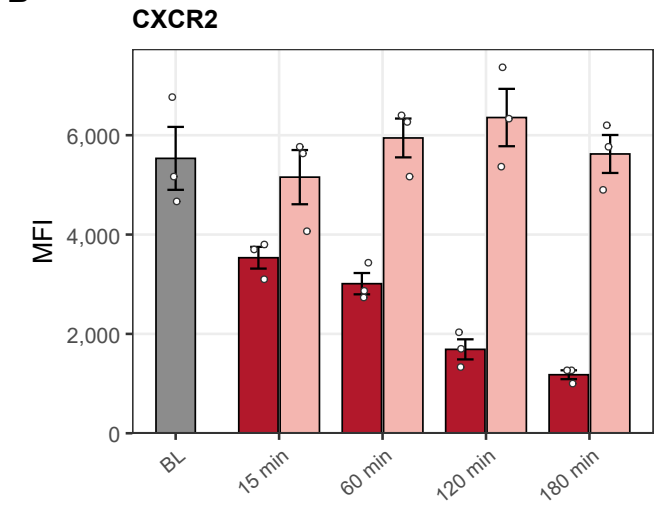

**C**

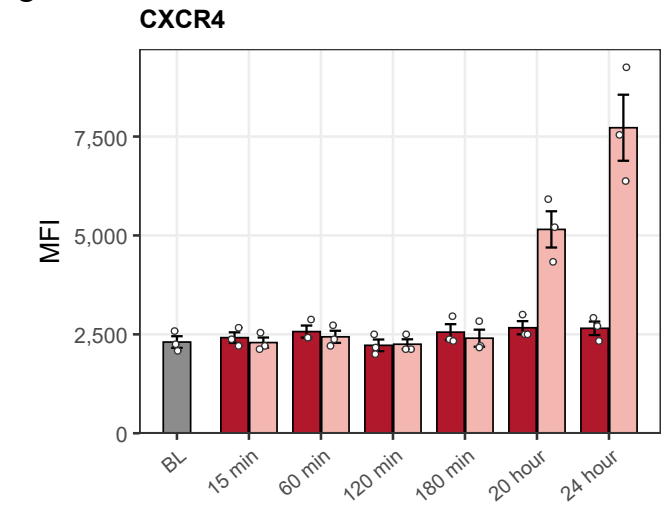

**D**

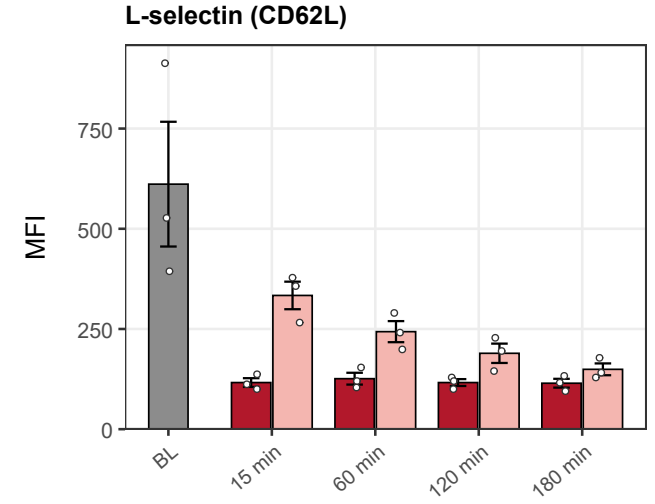

**E**

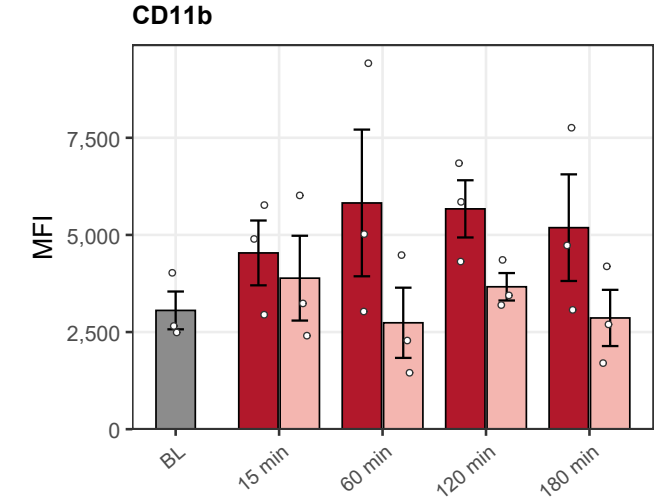

**F**

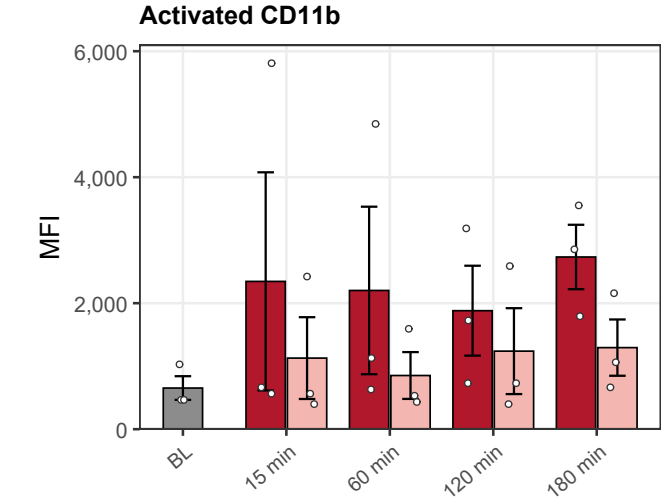

Baseline *S. aureus* Mock
